# Polyketide Synthase-Like Functionality Acquired by Plant Fatty Acid Elongase

**DOI:** 10.1101/2025.10.13.682109

**Authors:** Hyojin Kim, Kiyoul Park, Hao Jia, Fan Huang, Xinyu Tang, Ting Zhong, Wenlin Yu, Jean-Jack M. Riethoven, Rebecca E. Cahoon, Chunyu Zhang, Edgar B. Cahoon

## Abstract

Fatty acid elongation (FAE) typically proceeds through a four-step cycle of condensation, reduction, dehydration, and reduction to yield very long-chain fatty acids. Here, we describe a variation of this pathway in *Orychophragmus limprichtianus*, whose seed oil was found to contains previously unknown C24-C28 keto-hydroxy fatty acids that account for ∼25% of total fatty acids. Our studies revealed that these compounds are produced through an endoplasmic reticulum-localized discontinuous elongation process in which a 3-keto hydroxy intermediate bypasses full reduction and is extended through a polyketide synthase-like mechanism. Transcriptomic and functional assays identified two divergent enzymes, OlFAE1-2 and the low-activity ketoreductase OlKCR1-1, as central to this process. Protein modeling and mutant analysis suggest that specific amino acid substitutions underlie the altered activity of OlKCR1-1, enabling accumulation of keto intermediates. A portion of these unusual fatty acids accumulated in estolides, complex triacylglycerols that expand the chemical repertoire of seed oils. Together, our findings reveal unexpected flexibility in plant fatty acid elongation and provide new tools for engineering plants and microbes to produce renewable oils with tailored industrial functions.

## Introduction

The extraordinary chemical diversity of plant lipids represents a vast resource for sustainable bioproducts. Plant-derived triacylglycerols (TAGs), the primary components of vegetable oils, are crucial for nutrition and are increasingly engineered as feedstocks for biofuels and industrial applications, such as high-performance lubricants^1, 2, 3, 4^. The functionality of these oils is dictated by the unique structures of their constituent fatty acids, which can be modified to create new-to-nature molecules^5, 6^.

Plants have evolved diverse enzymatic strategies to produce novel fatty acids. While some modifications occur during the elongation cycle, a more common route involves tailoring enzymes that act on completed fatty acyl chains. These enzymes, including hydroxylases, desaturases, and epoxygenases, introduce functional groups such as hydroxyl, epoxy, and acetylenic bonds into the fatty acid backbone^7, 8, 9^. These post-synthesis modifications are responsible for many of the unusual fatty acids found in nature, such as the ricinoleic acid in castor bean oil^10^.

In contrast, other, rarer examples involve modifications of the core fatty acid elongation machinery itself (Fig. 1). One such example is the synthesis of very-long-chain fatty acids (VLCFAs) which typically occurs in the endoplasmic reticulum through a four-step cycle involving condensation, reduction, dehydration, and a final reduction^11^. We refer to this as the “continuous elongation” pathway, as it completes all four steps to extend the carbon chain without interruption^12^. In a related process, polyketide synthases (PKS) use similar chain-extension chemistry to produce a remarkable diversity of secondary metabolites, including antibiotics and flavonoids^13, 14^. A key distinction between these two pathways is that PKS can “skip” the reduction steps, leaving in-chain keto groups that expand the chemical and functional space of the products^14^.

**Figure 1.**
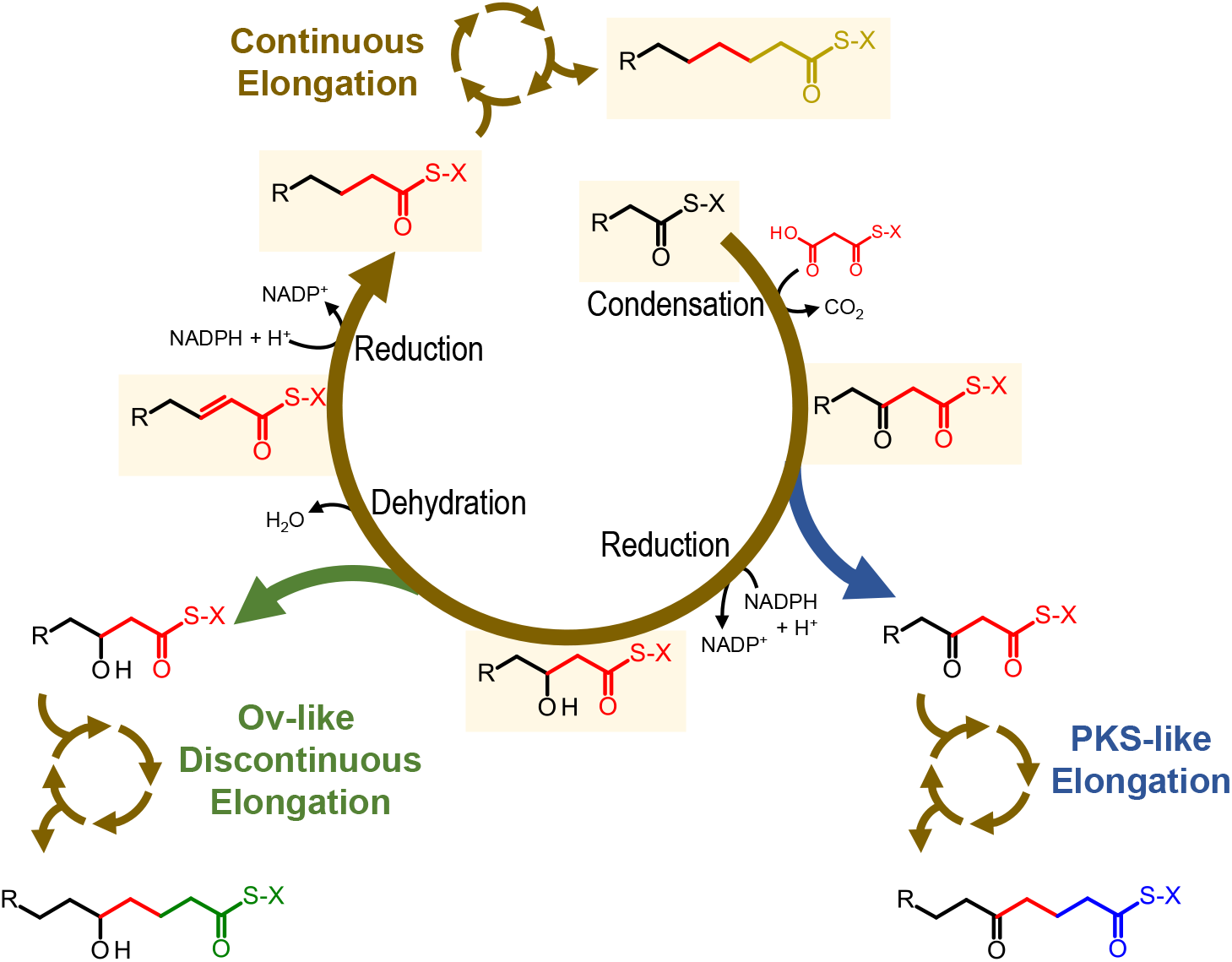
The simple pathway for fatty acid elongation in plant. One cycle of acyl chain elongation in the fatty acid elongation pathway is shown. Fatty acid elongation occurs through iterative cycles of acyl chain extension, involving condensation, keto-reduction, dehydration, and enol-reduction reactions. In-chain keto fatty acids are generated through polyketide (PKS)-like elongation of a 3-keto fatty acid intermediate. In-chain hydroxy fatty acids are synthesized through Ov-like discontinuous elongation of a 3-hydroxy fatty acid intermediate. X represents an acyl carrier protein (ACP) or coenzyme A (CoA). R represents a non-branched aliphatic carbon chain (= (CH_2_)n-CH_3_).

We previously discovered a “discontinuous elongation” pathway in *Orychophragmus violaceus* that deviates from the canonical VLCFA synthesis process by skipping the dehydration step^12^. This unique pathway, initiated by a variant fatty acid elongase 1 (FAE1), produces dihydroxy fatty acids whose TAGs form estolide-type oils with superior high-temperature lubricant properties with commercial potential. In this pathway, elongation of the C18 hydroxy fatty acid ricinoleic acid in its CoA form skips further elongation as 3-OH intermediate, prior to the dehydration step. This reaction, which is catalyzed by a variant FAE1, generates a C20 fatty acid with hydroxyl groups at its C3 and C20 positions. Two subsequent continuous elongation steps give rise to the C24 dihydroxy fatty acid nebraskanic acid and its omega-3 unsaturated variant wuhanic acid that accumulate to ≤40% of *O. violaceus* seed oil fatty acids. This finding revealed the inherent plasticity of plant fatty acid elongation and its potential for creating novel oil functionalities.

Here, we report the discovery of a new biosynthetic pathway in the seeds of *Orychophragmus limprichtianus*, a close relative of *O. violaceus*. We show that this species produces previously uncharacterized very-long-chain keto-hydroxy fatty acids in addition to dihydroxy fatty acids. We demonstrate that this unique composition arises from a hybrid metabolic pathway that combines discontinuous elongation with a polyketide synthase-like functionality. Our results identify two key divergent enzymes, a variant condensing enzyme (OlFAE1-2) and a reduced-activity ketoreductase (OlKCR1-1), that together form a novel enzymatic system for producing in-chain keto groups. We successfully reconstructed this pathway in an engineered oilseed host, demonstrating its potential for producing new-to-nature TAGs with expanded functionalities for high-performance bioproducts.

## Results

### Seed oil of O. limprichtianus contains keto-hydroxy fatty acids

Based on our previous work with the *Orychophragmus* genus, we initially characterized the fatty acid profile of *O. limprichtianus* seeds to determine if they contained unusual fatty acids. We compared extracts from *O. limprichtianus* (Ol) seeds with those from *O. violaceus* (Ov; didydroxy fatty acid-rich), castor bean (monohydroxy fatty acid-rich), and soybean (typical fatty acid-rich) (Fig. 2a-c) using thin-layer chromatography (TLC). These analyses revealed that Ol and Ov seeds contain little or no typical triacylglycerols (TAG). Instead, Ol seeds have a major, high mobility TAG band and a fainter band similar to TAG estolides that occur in Ov seeds^15^ (Fig. 2b). Fatty acid methyl esters (FAMEs) generated from Ol seeds also showed high-mobility bands not present in castor bean oil and distinct from Ov dihydroxy FAMEs (Fig. 2c).

**Figure 2.**
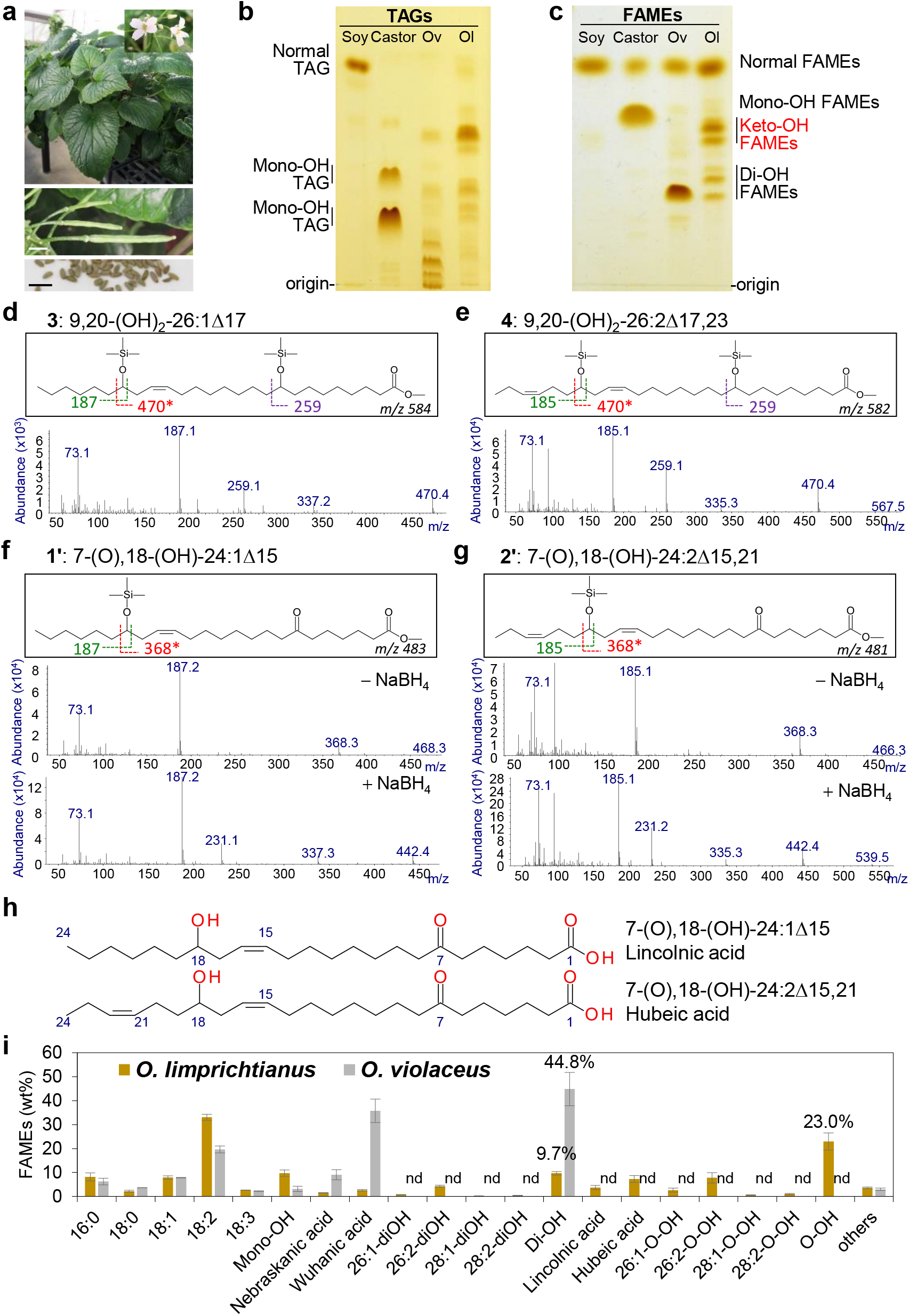
Identification of novel fatty acids in *O. limprichtianus* seeds. **a**, Images of *O. limprichtianus* (Ol) plant and seeds. Scale bar=5mm. b and c, Silica TLC Separation of oil (**b**) and FAMEs (**c**) from soybean, castor, Ov and Ol seeds. Total oils or FAMEs prepared from soybean, castor, Ov and Ol seeds were separated on TLC plate with mobile phase [Diethyl Ether: Heptane; 65:35]. Mono-OH, monohydroxy fatty acids. Di-OH, dihydroxy fatty acids. **d** to **g**, Identification of unusual hydroxy fatty acids using GC-MS analysis. Black boxes present fragment ion patterns of peak 3, 4, 1’, and 2’ from Supplementary Figure 2 and 3. The asterisks for ion ‘m/z=396’ and ‘m/z=470’ in black box are formed by a complex rearrangement involving transfer of the TMS group to the carboxyl group. **h**, Structures of lincolnic acid and hubeic acid. **i**, Fatty acid composition of two *Orychophragmus* seeds. Values are presented as wt% of total fatty acids. Di-OH, dihydroxy fatty acids. O-OH, keto-hydroxy fatty acids. Others include 20:0, 20:1, 20:2, 22:0, 22:1, 24:0, and 24:1 fatty acids. The results are the average of three replicates ± standard deviation.

To identify the molecular structures of the FAMEs separated on TLC, we performed gas chromatography (GC) and GC-mass spectrometry (MS) on the trimethylsilyl-derivatized FAMEs. This analysis revealed mono- and di-unsaturated C24 dihydroxy fatty acids (nebraskanic and wuhanic acids) as well as the C26, and C28 elongated forms of nebraskanic and wuhanic acids. We also discovered six major, previously uncharacterized fatty acids with longer retention times (Fig. 2d,e; Supplementary Figs. 1-4). The mass spectra of these fatty acids led us to hypothesize that they are keto-hydroxy fatty acids. To definitively test this hypothesis, we reacted the Ol FAMEs with sodium borohydride, which reduces keto groups to hydroxyl groups. This reaction specifically depleted the unknown fatty acid peaks and proportionally increased the peaks corresponding to dihydroxy fatty acids, including nebraskanic and wuhanic acids and their C26 and C28 elongated forms (Fig. 2f,g; Supplementary Figs. 1, 4). As shown in Fig. 2h, the fatty acids with the structures (7R,15Z,18R)-7-keto,18-hydroxy-tetracosa-15-enoic and (7R,15Z,18R,21Z)-7-keto,18-hydroxy-tetracosa-15,21-dienoic acids were designated as ‘lincolnic’ and ‘hubeic’ acids, respectively. Mass spectra of the other peaks were consistent with C26 9-keto, 20-hydroxy and C28 11-keto, 22-hydroxy mono- and di-unsaturated fatty acids (Supplementary Figs 2, 3). Subsequent quantification revealed that these novel keto-hydroxy fatty acids constitute 23% of the total fatty acids in the seed oil. (Fig. 1d). Overall, our analyses revealed that Ol TAG is distinct from that of Ov TAG by not only the presence of C26 and C28 dihydroxy fatty acids, but more notably by the occurrence of previously unreported C24-C28 keto-hydroxy fatty acids.

**Figure 3.**
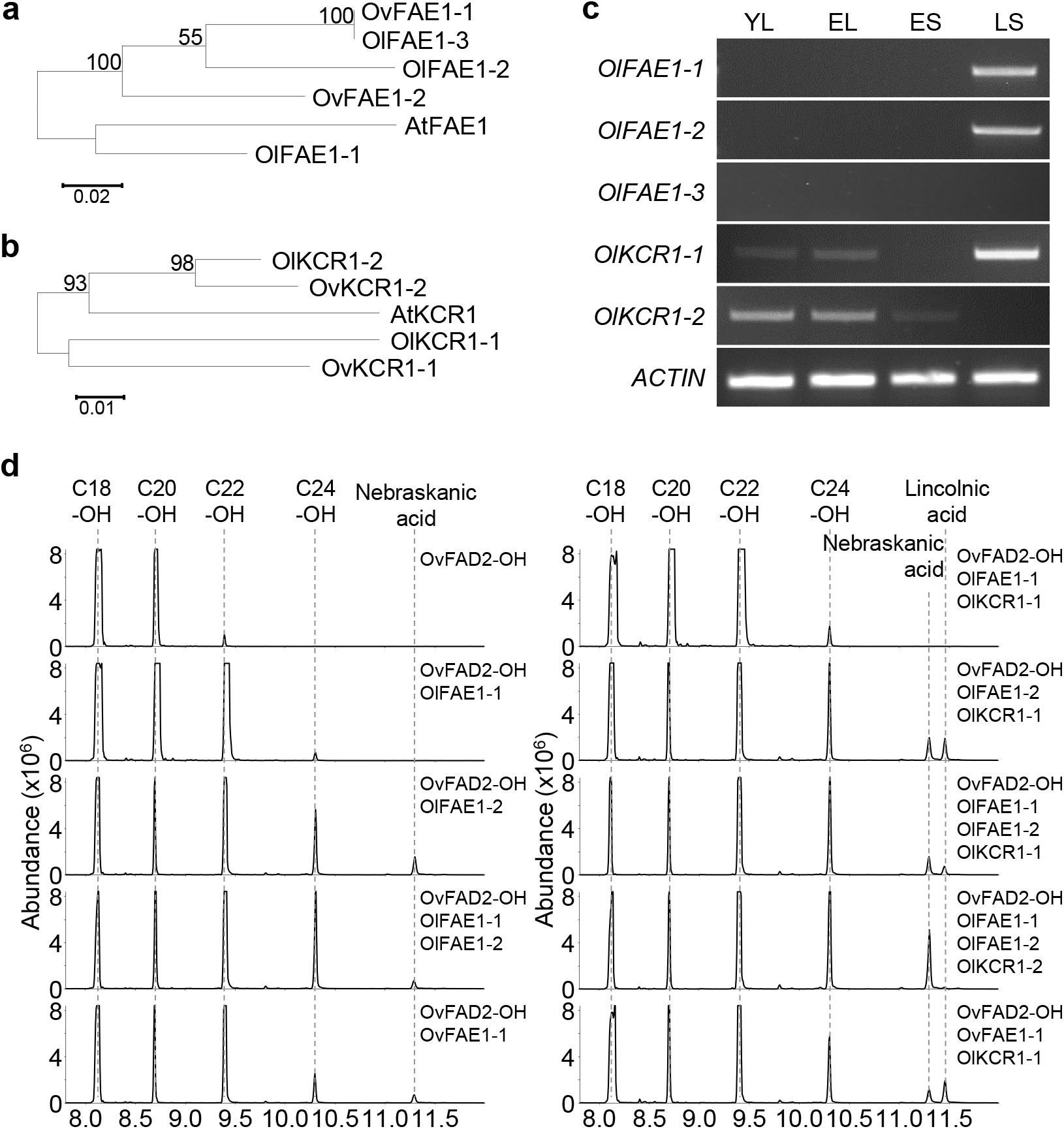
Functional analysis of *OlFAE1* and *OlKCR1* genes. **a** and **b**, Neighbor-joining phylogenic tree generated by clustalW alignment of deduced amino acid sequences of *FAE1* genes (a) or *KCR1* genes (b) from *A. thaliana, O. violaceus* and O. *limprichtianus* plants computed using MEGA 11. OvKCR1-1 and OvKCR1-2, OV09G031750 and OV12G016450 (http://www.bioinformaticslab.cn/pubs/OV_data/; Zhang et al. 2022). Numbers indicate branch support assessed by 500 bootstrap iterations (values below 50 are not shown). The tree is drawn to scale, with branch lengths in the same units as those of the evolutionary distances used to infer the phylogenetic tree. The evolutionary distances were computed using the JTT matrix-based method. **c**, RT-PCR analysis of three *OlFAE1* genes and two *OlKCR1* genes. cDNA from young leaf (YL), fully expanded leaf (EL), early developing seed (ES; before 30-days after flowering), and late developing seed (LS; after 30-day after flowering). **d**, GC-MS chromatograms of transgenic Col-0 plants. Total FAMEs from various transgenic Arabidopsis seeds were converted to TMS derivatives. Representive chromatograms showed the extracted ion m/z 187. NA, nebraskanic acid. LA, lincolnic acid.

**Figure 4.**
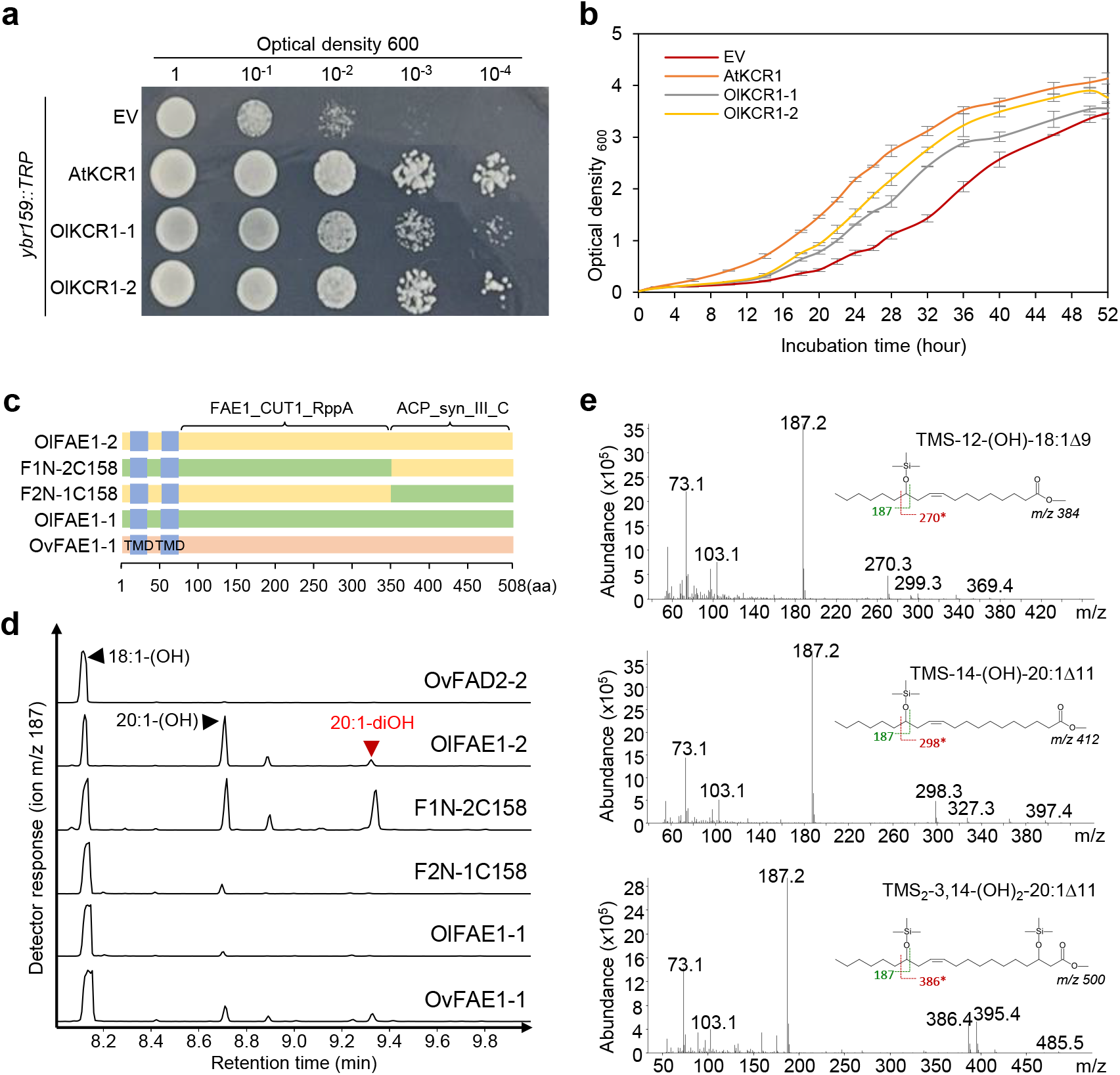
Domain analysis of FAE1 or KCR1 in yeast. **a**, Schematic view of native and chimeric FAE1 and KCR1 proteins. Two transmembrane domains (TMDs) possess at N-terminus of proteins. Domain swapped chimeric mutants were generated through overlapping PCR. F1N-2C158, OlFAE1-1^(1-350)^ + OlFAE1-2^(350-507)^. F2N-1C158, OlFAE1-2^(1-349)^ + OlFAE1-1^(351-508)^. K1N-2C21, OlKCR1-1^(1-297)^ + OlKCR1-2^(298-318)^. K2N-1C21, OlKCR1-2^(1-297)^ + OlKCR1-1^(298-318)^. **b**, GC-MS analysis after yeast induction assay using FAE1 proteins. GC-MS analyses of FAMEs prepared from 2-day induced yeast cell culture with trimethylsilyl derivatization of hydroxylated fatty acids The chromatograms shown are representative of analyses of greater than five independent biological samples, with each showing similar fatty acid profiles. **c**, Mass spectra of TMS-derivatives of 18:1-OH, 20:1-OH and 20:1-diOH. The asterisks for ion ‘m/z = 299, 327 or 386’ are formed by a complex rearrangement involving transfer of the TMS group to the carboxyl group. **d** and **e**, Growth complement assay in *ybr159* yeast mutant. Vectors were introduced into *ybr159::TRP* strain. Cells harboring designated vector were inoculated under selective solid and liquid media (SD-Leu-Trp) at 30°C.

### OlFAE1-2 and OlKCR1-1 are divergent enzymes for biosynthesis of in-chain keto groups

We subsequently conducted transcriptomic analysis of developing *O. limprichtianus* seeds to identify candidate genes responsible for the biosynthesis of the in-chain keto group. Our GC-MS analysis in this study, combined with our previous work^12^, supported a model where these unusual fatty acids are generated from specific C20 intermediates within the fatty acid elongation cycle. In particular, the C24 dihydroxy fatty acids in Ov seeds arise from a C20 3-hydroxy lesquerolic acid acyl-CoA intermediate produced by the combined activities of an ER-localized FAE1 and KCR1, followed by the skipping of the subsequent dehydration and reduction reactions in the fatty acid elongation^12^. In contrast, we hypothesized that the novel Ol C24 keto-hydroxy fatty acids are generated from a C20 3-keto lesquerolic acid intermediate, which is produced when FAE1 acts without a subsequent KCR1 reduction step. This essentially ends the elongation cycle prematurely, after the first of the four typical elongation reactions, to generate the C20 3-ketoacyl-CoA intermediate, which is subsequently elongated through two complete cycles to yield a C24 7-ketoacyl-CoA.

As a first step toward testing this model, we conducted transcriptomic analysis of developing *O. limprichtianus* (Ol) seeds and genomic DNA-based PCR analysis in leaves to identify candidate genes associated with the biosynthesis of the in-chain keto group. We compared the *FAE1* and *KCR1* genes from Ol, Ov, and other Brassicaceae plants, which showed a high degree of identity in both their nucleotide and deduced amino acid sequences (Supplementary Figs. 5, 6). The phylogenetic analysis revealed that OlFAE1-2 clusters together in the same group with OvFAE1-1 and OlFAE1-3, which have identical sequences. This suggests that OlFAE1-2 likely plays a role in dihydroxy fatty acid elongation, similar to OvFAE1-1. By contrast, OlFAE1-1 clustered most closely with the Arabidopsis FAE1 (AtFAE1), suggesting that this enzyme is a typical FAE1 that is associated with continuous elongation. In addition, the two OlKCR1 sequences separated into distinct clades, indicating potentially functional differences between these enzymes (Fig. 3a,b). Notably OlKCR1-1 clusters with OvKCR1-1, and OlKCR1-2 clusters with the Arabidopsis KCR (AtKCR1). Gene expression analysis confirmed that OlFAE1-1, OlFAE1-2, and OlKCR1-1 are expressed during late seed development in Ol plants, whereas expression of OlKCR1-2 is expressed primarily in leaves. In contrast, OlFAE1-3 exhibited pseudogene transcripts, indicating that it is a nonfunctional gene (Fig. 3c and Supplementary Fig. 7). Based on their divergence from the Arabidopsis homologs and their seed expression, OlFAE1-2 and OlKCR1-1 appeared to be the best candidates for association with keto-hydroxy fatty acid biosynthesis.

**Figure 5.**
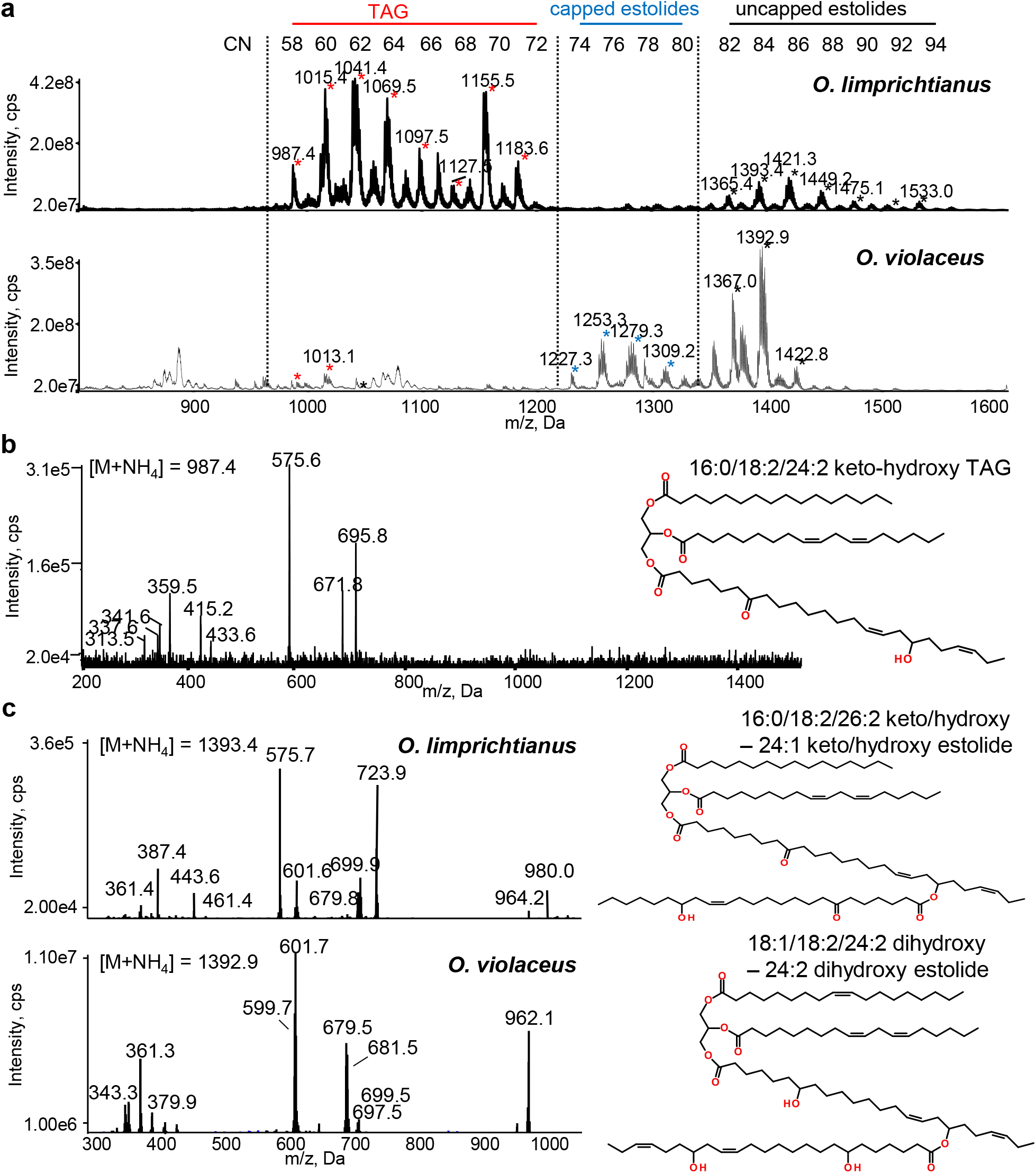
Ol estolide structure characterization. **a**, Q1 scan of total neutral lipid species extracted from Ol and Ov seeds. Starred peaks are M + NH_4_ TAG and estolide species that vary by carbon number (CN) as shown. In Ol samples, intermediate peaks represent mass differences due to loss of water (in-source fragmentation), and/or differences in hydroxylation or keto substitutions. **b**, Product ion spectrum and structure of the Ol 16:0/18:2/24:2 (7-O,18-OH) TAG (M + NH_4_), 986.8 m/z. The ions (m/z) detected were: 313.5 m/z = 16:0 R’ + 74. 337.6 m/z = 18:2 R’ + 74. 341.6 = 24:2 (7-O,18-OH) RCO+ - 2H_2_0. 359.5 m/z = 24:2 (7-O,18-OH) RCO+ - H_2_0. 415.2 m/z = 24:2 (7-O,18-OH) R’ + 74 –2H_2_0. 433.6 m/z = 24:2 (7-O,18-OH) R’ + 74 – H_2_0. 575.6 m/z = 16:0/18:2 DAG. 671.8 m/z = 16:0/24:2 (7-O,18-OH) DAG. 695.8 m/z = 18:2/24:2 (7-O,18-OH) DAG. **c**, Product ion spectra and representative structures of estolides detected in Ol (m/z 1393.4) and Ov (m/z 1392.9) seed oils. The spectra indicate the presence of two possible estolide species in each plant; the predominant structure for each is shown. Ol: 16:0/18:2/26:2 (7-O,18-OH)–24:1 (7-O,18-OH) estolide; Ov: 18:1/18:2/24:2 (7-,18-OH)–24:2 (7-,18-OH) estolide. Alternative isomers are presented in Supplementary Figure S14.

**Figure 6.**
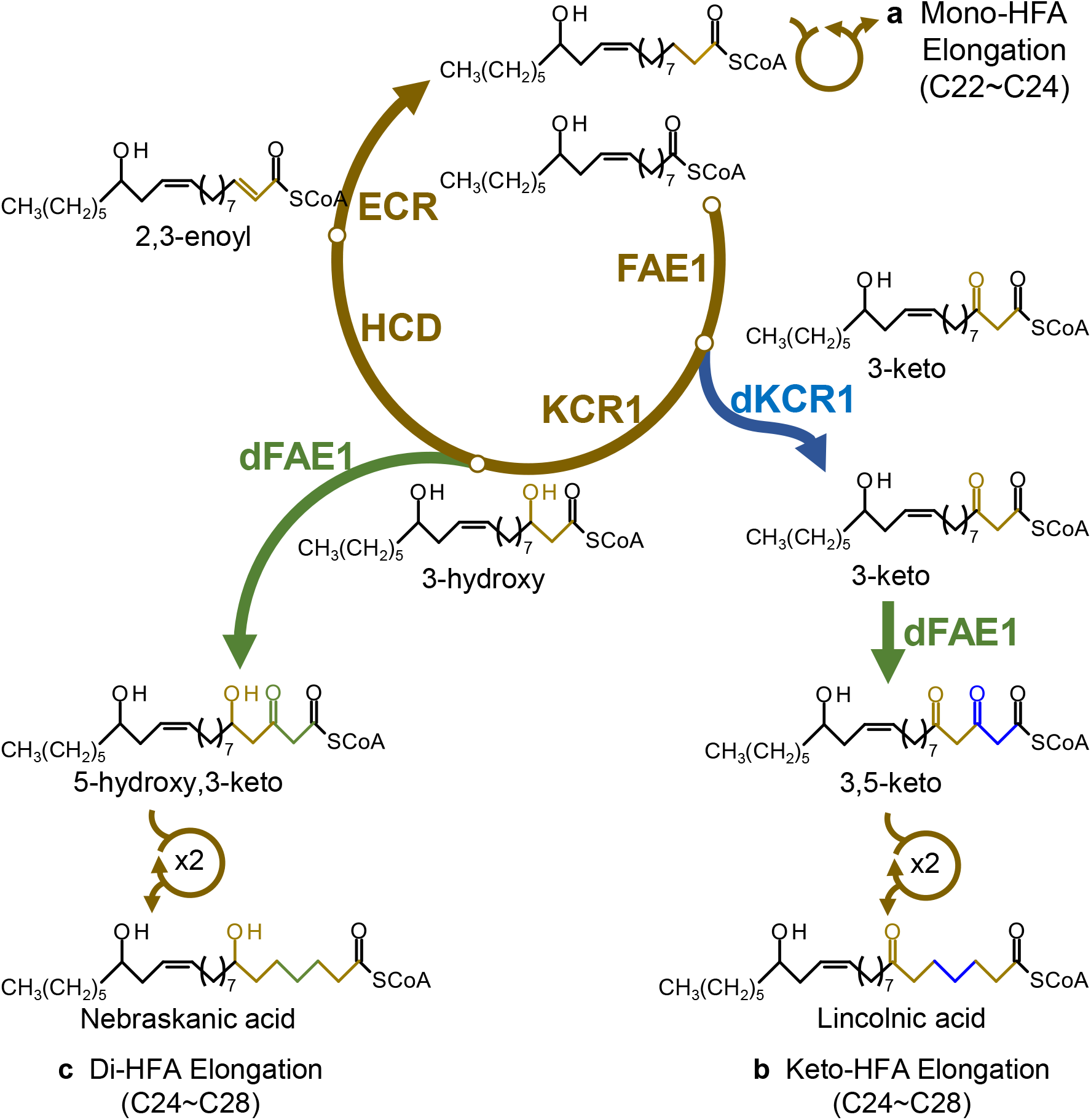
Elongation pathways of hydroxy fatty acids in *Orychophragmus* species. **a**, Mono-hydroxy fatty acyl-CoAs are processed through the canonical fatty acid elongation (FAE) complex, beginning with a condensation reaction between malonyl-CoA and ricinoleoyl-CoA, yielding chain-elongated products ranging from C20 to C24. **b**, Keto-hydroxy fatty acid biosynthesis proceeds via a PKS-like elongation pathway. Due to impaired 3-ketoacyl-CoA reductase (KCR) activity of the divergent KCR1 (dKCR1), the divergent FAE1 (dFAE1) intercepts 3-keto-hydroxy fatty acyl-CoAs and elongates them into 3,5-keto-hydroxy fatty acyl-CoAs, ultimately generating keto-hydroxy fatty acyl-CoAs with chain lengths from C24 to C28. **c**, Di-hydroxy fatty acyl-CoAs are elongated via an Ov-like discontinuous elongation pathway. In this process, dFAE1 catalyzes a condensation reaction between malonyl-CoA and a 3-hydroxy fatty acyl-CoA, producing di-hydroxy fatty acyl-CoAs ranging from C24 to C28. Brown arrows indicate the canonical continuous fatty acid elongation pathway. Green arrows represent the Ov-like discontinuous elongation pathway, wherein dFAE1 catalyzes the condensation of malonyl-CoA with 3-hydroxy acyl-CoA intermediates. The blue arrow indicates the PKS-like elongation pathway, in which elongation is interrupted due to reduced dKCR1 activity, allowing dFAE1 to act on 3-keto-hydroxy intermediates. FAE1, fatty acid elongase 1. dFAE1, divergent FAE1. KCR1, 3-ketoacyl CoA reductase 1. dKCR1, divergent KCR1. HCD, 3-hydroxy CoA dehydrogenase. ECR, 2,3-enoyl CoA reductase.

### Reconstitution of keto-hydroxy fatty acid biosynthesis in Arabidopsis

To determine the functions of the *OlFAE1s* and *OlKCR1s*, we created transgenic *Arabidopsis* lines expressing various combinations of these genes under control of seed-specific promoters (Supplementary Fig. 8). We also co-expressed the fatty acid hydroxylase *OvFAD2-2* to generate the C18 monohydroxy fatty acyl (ricinoleoyl)-CoA that we previously showed is the precursor of the dihydroxy fatty acid nebraskanic acid. We then analyzed the fatty acid composition of seed oil from the engineered plants using TLC and GC-MS to determine reaction products. Co-expression of *OlFAE1-2* or *OvFAE1-1* (positive control) with *OvFAD2-2* resulted in the synthesis of triacylglycerols (TAGs) containing the dihydroxy fatty acid nebraskanic acid, along with C20-C24 monohydroxy fatty acids. By contrast, co-expression of *OlFAE1-1* and *OvFAD2-2* yielded only C20-C24 monohydroxy fatty acids. These results indicate that OlFAE1-2, like OvFAE1-1, can initiate discontinuous as well as continuous elongation, whereas OlFAE1-1 only initiates continuous elongation. The introduction of 3-ketoacyl-CoA reductase (*OlKCR1-1*) resulted in the production of nebraskanic acid along with the keto-hydroxy fatty acid lincolnic acid to amounts of <3% of the total oil (Supplementary Figs. 9-11). This result was also replicated by expression of *OvFAE1-1* with *OlKCR1-1*. Using OlKCR1-2, which has more typical sequence, we detected only nebraskanic acid, but no lincolnic acid. Notably, the gene combination with *OlFAE1-1, OlFAE1-2*, and *OlKCR1-1* yielded nebraskanic acid levels of <5% of the total oil (Supplementary Figs. 11b). Overall, these results confirm that OlFAE1-2, as well as OvFAE1-1, in combination with OlKCR1-1 generates the in-chain keto group found in lincolnic acid. This is consistent with acquisition of PKS functionality in plant fatty acid elongation that uses the CoA ester of 3-keto lesquerolic acid intermediate, with subsequent continuous elongation, to generate lincolnic and other keto-hydroxy fatty acids in Ol seeds.

### Reduced activity of OlKCR1-1 revealed by mutant analysis

The results above indicate that expression of *OlKCR1-1*, rather than *OlKCR1-2*, in combination with divergent *FAE1s* from Ol or Ov results in production of keto-hydroxy fatty acids. We hypothesized that OlKCR1-1 has slower catalytic turnover than OlKCR1-2, which would allow the divergent FAE1 enzymes access to higher pools of the 3-keto intermediate for PKS-like functionality. To test this hypothesis, we compared the ability of OlKCR1-1 and OlKCR1-2 to complement the *Saccharomyces cerevisiae ybr159::Trp1* mutant. This mutant is deficient in 3-ketoreductase activity that results in a deficiency of very-long chain fatty acids, leading to a slow-growth phenotype^16^. For these complementation assays, we also used the AtKCR1 as a positive control. We found that the introduction of AtKCR1, OlKCR1-1 and OlKCR1-2 significantly improved the growth rate of the 3-ketoreductase mutant relative to an empty vector control. However, the growth rate observed with the introduction of OlKCR1-1 was lower than that with AtKCR1 or OlKCR1-2 (Fig. 4a,b), consistent with a lower activity of OlKCR1-1.

We next examined the structural basis for the reduced activity of OlKCR1-1 versus OlKCR1-2. These enzymes share 89% amino acid identity, including the highly conserved the NADH-binding motif (GX_3_GXGX_3_AX_3_AX_2_G, positions 58-75) and the essential catalytic motif (S-Y-K; positions 191, 206, and 210)^17^ (Supplementary Fig. 6b). Using AlphaFold3 to examine the three-dimensional structures of OlKCR1-1 and OlKCR1-2^18^, we found that the predicted template modeling (pTM) score and the interface predicted template modeling (ipTM) score of each KCR1 protein were above 0.5 and 0.8 (AtKCR1, 0.93 and 0.95; OlKCR1-1, 0.92 and 0.95; OlKCR1-2, 0.93 and 0.95), respectively. This indicated that the AlphaFold 3 predictions for the KCR1 proteins are highly confident and of high quality. The analysis revealed that the C-terminal helix is positioned in close proximity to the FA-binding pocket (Supplementary Fig.12). Since FA binding is influenced by the surrounding electrostatic environment, R301 likely plays a critical role in substrate-enzyme interactions. This residue is highly conserved among plant KCR enzymes (Supplementary Fig. 6b). However, in the divergent OlKCR1-1, R301 is substituted with serine (R301S). Based on AlphaFold 3 predictions, R301S likely not only neutralizes the electrostatic surface potential in this region but also alters the geometry of the FA-binding pocket (Supplementary Fig. 12b). This substitution may impair the normal substrate-enzyme interaction and catalysis in OlKCR1-1, potentially leading to altered enzyme activity.

Based on this modeling, we then swapped 21 amino acids of the C-terminus of OlKCR1-2 into the corresponding region of OlKCR1-1. This region included the amino acid difference at residue 301, as well as three other amino acid differences between OlKCR1-1 and -2. This chimeric enzyme enhanced *ybr159* growth, similar to that conferred by OlKCR1-2 (Supplementary Fig. 13). These results indicate that amino acid differences in the OlKCR1-1 C-terminus contributes to reduced 3-ketoreductase activity. Notably, expression of the reciprocal swap of the C-terminal amino acids of OlKCR1-1 in place of the corresponding region of OlKCR1-2 did not result in observable growth reduction relative to the OlKCR1-2 in the *ybr* mutant, suggesting that additional amino acid differences in OlKCR1-1 contribute to the apparent lower activity of this enzyme.

### Discontinuous FAE1 elongation

We also examined the regions of FAE1 that are associated with the divergent activity of OlFAE1-2. For these experiments, we used *S. cerevisiae* as a model system and co-expressed OvFAD2-2 hydroxylase to generate the ricinoyl-CoA (12OH-18:1-CoA) substrate of OlFAE1-2. Due to the high identity between the typical OlFAE1-1 and divergent OlFAE1-2 (Supplementary Fig. 5), we designed the domain swapped structures based on the functional domains: N-region including a domain for FAE1/Type III polyketide synthase-like protein which also are required for interaction with other FAE components and elongation activity, and C-region including a domain for 3-oxoacyl-(ACP) synthase III^19, 20, 21^ (Fig 4c). The co-expression of *OlFAE1-2* and *OvFAD2-2* synthesized both mono-HFAs (18:1-OH and 20:1-OH) and C20 3-hydroxy-lesquerolic acid (20:1-(OH)_2_), the product of the initial discontinuous elongation cycle. In the yeast system, we did not detect further elongation of the C20 dihydroxy fatty acid. In contrast, co-expression of the typical *OlFAE1-1* and *OvFAD2-2* synthesized only mono-HFAs (18:1-OH and 20:1-OH) (Fig. 4d,e). Co-expression of *OvFAD2-2* and OlFAE1-1 chimera (F1N-2C158) containing C-region of OlFAE1-2 exhibited the production of the C20 3-hydroxy-lesquerolic acid (Fig. 4d,e). However, the converse swap of the N-terminal region of OlFAE1-2 in place of the corresponding portion of OlFAE1-1 did not confer the ability of the chimeric enzyme to generate dihydroxy fatty acids. These results indicate that the C-terminal region of OlFAE1 plays a critical role in determining whether the enzyme exhibits continuous or discontinuous elongation activity.

### Estolides in O. limprichtianus seed oil

The seed oil of *O. limprichtianus* contains various hydroxy fatty acids, including mono-, di-, and keto-hydroxy species, whose chemical structures are well suited to form estolides, similar to those reported in *O. violaceus* seed oil. TLC analysis revealed a distinct TAG separation pattern (Fig. 2b), which prompted further LC-MS analysis to determine whether oligomeric estolides are naturally present in the *O. limprichtianus* seed oil. The majority of lipid species in *O. limprichtianus* seed oil were identified as typical TAGs (m/z 987-1183), while uncapped estolides were also detected across a broad m/z range (m/z 1365-1533) (Fig. 5a). In contrast, the *O. violaceus* seed oil, included here as a reference, was composed almost exclusively of estolides with negligible amounts of typical TAGs, consistent with the findings of Romsdahl et al. (2019) (Fig. 5a).

LC-MS/MS product ion analysis further confirmed the incorporation of keto-hydroxy fatty acids into TAG molecules. A representative spectrum of the precursor ion at m/z 987.4 corresponded to a TAG species containing 16:0, 18:2, and 24:2 (7-O, 18-OH) (Fig. 5b). Moreover, uncapped estolides in *O. limprichtianus* were observed over a broader m/z range (1365-1533) compared to those in *O. violaceus* (Fig. 5a). For instance, the ion at m/z 1393 in *O. limprichtianus* was identified as an estolide consisting of TAG (16:0/18:2/26:2 (7-O,18-OH)) esterified with 24:1 (7-O,18-OH), whereas the corresponding species in *O. violaceus* was composed of a TAG backbone 18:1, 18:2, and 24:2 (7-,18-OH) esterified with another 24:2 (7-,18-OH) chain (Fig. 5c). In addition, alternative estolide structures with the same precursor ion (m/z 1393) were inferred from product ion spectra: 18:1/18:1/24:2 keto/hydroxy–24:1 keto/hydroxy estolide in *O. limprichtianus* and 18:2/18:2/24:1 dihydroxy– 24:2 dihydroxy estolide in *O. violaceus* (Fig. S14). Taken together, these results demonstrate that *O. limprichtianus* seed oil consists primarily of typical TAGs along with diverse uncapped estolides containing hydroxy and keto-hydroxy fatty acids, in clear contrast to the predominantly estolide-based composition of *O. violaceus* seed oil.

## Discussion

In this study, we identified previously unknown C24-C28 keto-hydroxy fatty acids as major components of *O. limprichtianus* seed oil. These unusual structures, including the C24 species lincolnic and hubeic acids, differ from the corresponding dihydroxy fatty acids of *O. violaceus* by the substitution of the mid-chain hydroxyl (e.g., 7-hydroxy in C24, 9-hydroxy in C26) with a keto group. The discovery of these new fatty acids prompted us to investigate their biosynthetic origin, leading to the elucidation of an ER-localized elongation pathway that diverges from the canonical fatty acid elongation cycle and functionally resembles polyketide synthases. We show that this pathway arises from the combined activity of a divergent FAE1 (OlFAE1-2), capable of condensing 3-hydroxy or 3-keto intermediates, and a less active KCR isoform (OlKCR1-1), which allows the 3-keto intermediate to persist in the elongated fatty acid chains. Together, these enzymes generate keto-hydroxy fatty acids through a modified form of discontinuous elongation, demonstrating remarkable plasticity in ER-associated fatty acid metabolism. The outcomes of these reactions yield novel fatty acid structures with enhanced functionality for bioproduct applications.

While this study focused on metabolic diversity discovery, the mechanisms underlying the enzymes associated with discontinuous fatty acid elongation remain to be determined. In particular, the structural basis that enables the divergent FAE1 to catalyze condensation using 3-hydroxy or 3-keto intermediates and to catalyze this reaction specially at the C18 to C20 elongation stage is not clear. In addition, based on fatty acid composition of *O. limprichtianus* seed oil and Arabidopsis metabolic engineering, it appears that this pathway is initiated by elongation of the CoA ester of a C18 12-monohydroxy fatty acid (ricinoleic acid), and the variant FAE uses the 3-keto C20 intermediate as a condensation substrate. Through domain swapping, we identified the C’-region including a domain for 3-oxoacyl-(ACP) synthase III^19, 20, 21^ as critical for discontinuous elongation. In support of this, swapping of this domain into the typical *O. limprichtianus* FAE1-1 (F1N-2C158 in Fig. 4) conferred discontinuous elongation. Notably, differences in this catalytic domain across ketoacyl-CoA/ACP synthases is associated with the wide substrate specificity of these enzymes as part of fatty acid synthases and polyketide synthases found in nature to generate diverse chemical structures. In the case of the typical and divergent *O. limprichtianus* FAE1s, we presume that amino acid differences in this domain result in variations in the active site geometry to accommodate alternative substrate structures. However, our cursory comparative modeling using AlphaFold failed to provide mechanistic clues from the predicted structures of these enzymes. As we continue to explore the discontinuous elongation mechanism, structural data for FAE1 enzymes, which is currently limiting, will be critical. Of particular emphasis for these studies are comparisons among Brassicaceae FAE1s, including the *Physaria fendleri* FAE1 that has substrate preference for CoA esters of C18 12-monohydroxy fatty acids.

Our studies also highlighted other aspects of variant fatty acid elongation. Notably, we show that production of in-chain keto group requires expression of a divergent 3-ketoacyl-CoA reductase OlKCR1-1 in conjunction with the divergent FAE1. The most distinguishing feature of the primary structure of OlKCR1-1 is R301S substitution in this enzyme relative to KCR1 enzymes in other Brassicaceae. This substitution is predicted to affect substrate binding. Consistent with this, OlKCR1-1 expression in a yeast KCR mutant has less ability to complement the reduced growth rate of this mutant compared to expression of OlKCR1-2 and AtKCR1. As the most highly expressed KCR in *O. limprichtianus* seeds, OlKCR1-1 likely retains sufficient activity to support essential fatty acid elongation for pathways such as sphingolipid biosynthesis. We are currently exploring the hypothesis that any lower activity KCR functioning together with OlFAE1-2 or OvFAE1-1 can confer in-chain keto production. Of note, the ER-based fatty acid elongation system in plants has been proposed to function as a complex of the four enzymes required each continuous elongation step^11^. Given this possibility, we can also not rule out that the OlKCR1-1 has reduced affinity for interacting FAE1 enzymes, leading to transitory increase in 3-keto substrate for discontinuous elongation.

Another distinctive feature of *O. limprichtianus* oil composition is the enrichment of C26 and C28 keto-hydroxy and dihydroxy fatty acids, which are present in only low to non-detectable concentrations in *O. violaceus* seed oil. In contrast, C24 dihydroxy fatty acids are prevalent in *O. violaceus* seed oil^12^, and engineered Arabidopsis seeds accumulate primarily C24 fatty acids from discontinuous elongation. In general, the occurrence of C26 and C28 fatty acids is uncommon in seed oils of Brassicaceae species^22, 23^. Though not examined in the current study, these fatty acids may arise from association of the elongation system with a CER2 BAHD acyltransferase-like protein that has been shown to confer >C28 acyl chains in Arabidopsis wax biosynthesis^24^. Alternatively, a member of the KCS enzyme family with substrate specificity for C24 and C26 acyl-CoA substrates may be co-opted for the biosynthesis of *O. limprichtianus* seed oil fatty acids^11, 25^.

We previously showed that the TAG component of *O. violaceus* seed oil occurs almost completely in an estolide form^15^. These structures are comprised of a core TAG molecule with ≤eight additional fatty acids esterified to the hydroxyl group proximal to the methyl end of nebraskanic or wuhanic acids. Although TAG estolides occur in other hydroxy fatty acid-rich seed oils, such as those of the Brassicaceae *Physaria fendleri, O. violaceus* has the highest relative levels reported to date^12, 15, 26, 27, 28, 29, 30^. These TAG structures, whose synthesis is enabled by fatty acid hydroxylation, confer superior high-temperature lubricant functionality^15, 31, 32, 33, 34^. While lubricant performance of *O. violaceus* oil has been evaluated, comparable tests with *O. limprichtianus* oil have not been possible due to insufficient seed material. Our analysis nevertheless revealed TAG estolides in *O. limprichtianus* seed oil that contained keto-hydroxy as well as dihydroxy fatty acids. However, the relative concentrations of TAG estolides were lower than found in *O. violaceus* seeds. We also failed to detect estolides in the seed oil of Arabidopsis that produce dihydroxy fatty acids. These findings suggest that *O. violaceus* has evolved acyltransferase activity that is deficient in *O. limprichtianus* seeds and capable of generating high TAG estolide levels by acylation of the fatty acid methyl-terminal hydroxyl group.

Our studies to date with *O. limprichtianus* and *O. violaceus* seed oil has not only provided insights into the plasticity of fatty acid elongation for novel product outcomes but has also provided biotechnological tools for generating potential new high-value bioproducts in plant and microbial hosts. For example, we produced very long-chain dihydroxy fatty acids in Arabidopsis seeds using the divergent genes *OlFAE1-2* and *OvFAE1-1*, and generated very long-chain keto-hydroxy fatty acids when these genes were combined with *OlKCR1-1*, providing proof-of-principle for their biotechnological utility. Additional knowledge is needed about the elongation systems in *O. limprichtianus* and *O. violaceus*, including interactions among enzymes in this pathway, to maximize discontinuous elongation activity in heterologous hosts. We are also actively studying *O. violaceus* TAG estolide synthesis for biotechnological production of these novel fatty acid storage forms that enhance vegetable oil viscosity for biobased lubricant applications. While we are exploring development of *O. limprichtianus* and *O. violaceus* as new oilseed crops, biotechnological efforts to transfer fatty acid elongation and storage mechanisms from these plants to established crop and microbial hosts will ultimately enable production of oils with useful and tailored functionalities that are not currently found in nature.

## Methods

### Plant materials and growth conditions

*O. limprichtianus, O. violaceus* and *Arabidopsis thaliana* Col-0 and *fae1fad2* were grown at 25 °C with 13 h light/11 h dark in the greenhouse and vernalized at 10 °C with 10 h light/14 h dark for three weeks to induce flowering in the cold chamber. Arabidopsis plants were grown at 22 °C with 16 h light (120 μE m^−2^ s^−1^)/8 h dark in the controlled environment chamber.

### TLC analysis

Total lipids were extracted from ∼20 mg of Orychophragmus seeds, soybean seeds, and Arabidopsis seeds by a modified version of Bligh-Dyer method^35, 36^. FAMEs were prepared from Orychophragmus seeds, castor oil, Arabidopsis seeds, or *S. cerevisiae* cell pellets by transesterification in 2.5% sulfuric acid in methanol (v/v) as described^36^. Then, FAMEs were extracted using heptane. The extracted TAGs and FAMEs were separated by TLC on Silica 60 plates (10 × 20 cm, 0.25-mm layer thickness; Merck) using a solvent system of diethyl ether/heptane at a ratio of 65/35 (v/v) were exposed to iodine vapor in a sealed tank.

### GC analysis

The extracted FAMEs were dried and TMS-derivatized by adding 50 µl of pyridine and 50 µl of bis(trimethylsilyl)trifluoroacetamide (BSTFA)/trimethylchlorosilane (99/1, v/v) (Sigma-Aldrich), then mixtures were incubated at 90°C for 30 min. Dried FAMEs were reduced by adding 750 µl of NaBH_4_ solution (2.0 M in triethylene glycol dimethyl ether, Sigma-Aldrich), then mixtures incubated at 4°C for 16 hours, and then separated by adding 1 ml of 0.9% NaCl solution (in water) and 1 ml of heptane. Heptane extracts were dried under a nitrogen stream and TMS-derivatized.

TMS-derivatized FAMEs (with/without NaBH_4_) in heptane were analyzed using an Agilent 7890 GC with flame ionization detection (FID) or mass-selective detection (HP5973; Agilent) operating at an electron impact ionization potential of 70 eV. GC-FID analysis was resolved on a HP-INNOWax column (30 m × 0.25 mm, 0.25-µm film thickness; Agilent) and hydrogen was used as the carrier gas at an inlet flow rate of 66 ml min^−1^. The oven temperature was programmed: 200°C for 1min, then 200 to 250°C at 15°C min^-1^, hold 250°C for 15min; the inlet and FID temperatures were 350°C. GC-MS equipped with an HP-5 column (30 m × 0.32 mm, 0.25-µm film thickness; Agilent) operated with helium as carrier gas at an inlet flow rate of 8.5 ml min^-1^. The oven temperature was programmed: 90°C for 1 min, then up to 320°C at 30°C min^-1^, hold 320°C for 20 min; the inlet temperature was 250°C.

### RNA extraction and RNA-seq analysis

Total RNA was isolated from late-developing seeds of *O. limprichtianus* using the RNeasy Plant Mini Kit (Qiagen), following the manufacturer’s protocol. The extracted total RNA was treated with DNase I to remove contaminating genomic DNA.

Subsequently, the RNA was submitted for RNA sequencing (RNA-seq) (GENEWIZ) on a Pacific Biosciences (PacBio) Sequel II System (Pacific Biosciences. SMRT Link v10.1). The Iso-Seq protocol with HiFi (Circular Consensus Sequencing, CCS) chemistry was utilized for sequencing, and the adapters used were NEB_5p and NEB_Clontech_3p (Wenger et al. 2019). Bioinformatically, sequencing adapters were removed using Lima (v2.7.1) (https://github.com/PacificBiosciences/barcoding) with Iso-Seq settings (`--isoseq, -- peek_genes`), followed by refinement using `isoseq refine` (4.0.0) (https://github.com/PacificBiosciences/IsoSeq) to determine full-length non-chimeric (FLNC) reads and remove concatemers, while requiring the presence of a poly(A) tail (`--require-polya`). High-quality isoforms were generated using `isoseq cluster` with quality value support (`--use-qvs`) to collapse redundant FLNC reads into non-redundant, high-confidence transcript clusters. Annotation of the resulting transcripts was performed using a reciprocal BLAST^37, 38^ against the *A. thaliana* protein database (AraPort11)^39^. An in-house Python script was used to scan both BLAST output files to annotate the transcript based on reciprocal hits. These high-confidence, annotated transcripts were subsequently utilized to specifically identify orthologs of key fatty acid elongation genes in *O. limprichtianus*. Specifically, the sequences of AtFAE1 (At4G34520), OvFAE1-1 (KU187971.1), and AtKCR1 (At1G67730) served as bait sequences to query the transcriptome and identify OlFAE1 and OlKCR1 homologous genes.

### Gene isolation and Arabidopsis transformation

Since FAE1 is a well-known gene that lacks introns^40, 41, 42, 43^, we amplified the *FAE1* genomic DNA fragments from *O. limprichtianus* plant leaves using specific primers (FAE1_F and FAE1_R, as listed in Supplementary Table 1), which were designed based on the Arabidopsis *FAE1* gene nucleotide sequence to isolate the *FAE1* gene in *O. limprichtianus* plant. For isolation of *KCR1* gene sequence, gene-specific primers were designed based on the *OvKCR1* genome sequence (KCR1_F and KCR1_R, as listed in Supplementary Table 1) and used to amplify the *OlKCR1* gene from leaf cDNA and developing seed cDNA of *O. limprichtianus*.

The binary vector harboring the fatty acid desaturase/hydroxylase (OvFAD2) used in Li et al. (2018), hereafter referred to as ‘FD’, was modified to insert various *FAE1* and *KCR1* genes. The amplified DNA fragments with *Eco*RI (5’-end) and *Xho*I (3’-end) restriction sites of each *FAE1* and *KCR1* genes were digested and subcloned into pBinGlyRed2 binary vector using *Eco*RI and *Xho*I restriction sites. Gene cassette (Glycinin promoter::FAE1 or KCR1:Glycinin terminator) was digested using *Av*RII and *Spe*I restriction enzymes and stacked gene cassettes into ‘FD’ using *Spe*I restriction site to generate various binary vectors shown in Supplementary Figure 8. Stable transgenic Arabidopsis plants were generated using Agrobacterium-mediated floral dip method^44^. After transformation, transgenic seeds were screened for DsRed fluorescence signal. T2 seeds were tested for further experiments.

### Yeast FAE1 induction assay

Yeast expression vector, pESC-His (Agilent), was used to induce both OvFAD2-2 and variant FAE1 in *S. cerevisiae* BY4741 strain. OvFAD2-2 DNA fragment from FD binary vector and ligated into pESC1-HIS vector using *Eco*RI and *Not*I restriction sites. Each FAE1 DNA fragment was tagged with *Eco*RI (5’-end) and *Xho*I (3’-end) restriction sites, and subcloned into pYES2 vector. FAE1 DNA fragments were inserted out from pYES2 vector using *Bam*HI and *Xho*I restriction enzymes and ligated into pESC1-HIS + OvFAD2-2 with *Bam*HI and *Sal*I sites. Four Chimeric FAE1s were generated by overlapping PCR and subcloned as described above. Vector constructs were transformed into BY4741 strain using the PEG-mediated transformation method (Zymo Research). Selected positive colonies were further inoculated with liquid SD-His media with 2% glucose and transgenes were induced by media change to SD-His growth media with 2% raffinose and galactose. After 48 h cultivation, cells were harvested by centrifuge and total fatty acids were extracted and analyzed.

### Yeast growth complementation assay

The *AtKCR1* gene (At1G67730) was amplified from *A. thaliana* genomic DNA using PCR and used as a control for normal KCR1 activity. Each coding sequence (CDS) of KCR1s was tagged with *Eco*RI (5’-end) and *Xho*I (3’-end) restriction sites, then subcloned into the yeast expression vector pYX242 for constitutive expression. Two chimeric KCR1s were generated by overlapping PCR and subcloned as described above. Vector constructs were transformed into *ybr159* mutant strain (*ybr159::Trp1*) using the PEG-mediated transformation method and plated on selective media (SD-Leu-Trp). Yeast cells were further inoculated in liquid YPD media for 2 days at 30°C, washed three times with distilled water (ddH_2_O), and OD_600_ was measured using Beckman Coulter DU800 spectrophotometer. After that, cells were subsequently dotted onto selective plates and grown for 3 days at 30°C. For growth curve analysis, yeast cultures were grown in 100 ml of SD-Leu-TRP liquid media at 30°C with shaking at 180 rpm, with OD_600_ measurements taken every 2 hours.

### Estolide analysis

Total neutral lipids were extracted from seeds of *O. limprichtianus* and *O. violaceus* using the Bligh-Dyer method^35^. The dried lipid extracts were resuspended in Methanol:Isopropanol:10 mM ammonium formate (94:5:1, v/v/v) and analyzed by direct infusion mass spectrometry on a Sciex QTRAP 4000 mass spectrometer. Initial lipid profiling was performed using a Q1 scan over the mass range of m/z 300 to 2200, with a scan time of 3 seconds per scan and an infusion rate of 0.65 ml hr^-1^. Instrument source parameters were maintained at a source temperature of 300°C, Declustering Potential (DP) of 90 V, and Entrance Potential (EP) of 10 V. TAGs and estolide species were identified based on their M+NH_4_ ions. To confirm the structure of potential estolide-containing TAGs, product ion scans (MS/MS) were performed on prominent M+NH_4_ ions to further elucidate their structure. As a representative example, the M+NH_4_ ion at m/z 986.8 (corresponding to the 16:0/18:2/24:2 keto/hydroxy TAG in *O. limprichtianus*) was selected for fragmentation. For fragmentation, a Collision Energy (CE) of 54 V and a Collision Exit Potential (CXP) of 11 V were applied. Key characteristic fragment ions, including R′ +74 fragments (e.g., m/z 313.5 for 16:0 and m/z 337.6 for 18:2) and diacylglycerol (DAG) loss fragments, were used to confirm the specific fatty acyl composition and the presence of the 24:2 keto/hydroxy chain, validating the TAG structure. Similar product ion scans were performed on M+NH_4_ intact estolide species,

## Data availability

Sequence data for OlFAE1-1, OlFAE1-2, OlFAE1-3, OlKCR1-1 and OlKCR1-2 have been submitted to GenBank (submission ID 3012412; submitted October 7, 2025). Accession numbers are pending due to the U.S. federal government shutdown and will be provided once available.

## Acknowledgement

We Thank to Dr. Teresa Dunn, Uniformed Services University of the Health Sciences, for providing *ybr159::Trp1* yeast strain. We dedicate this manuscript to the memory of John Ohlrogge for his inspiration to study novel plant fatty acids.

Pathway engineering research was supported by the DOE Center for Advanced Bioenergy and Bioproducts Innovation funded U.S. Department of Energy (DOE), Office of Science, Biological and Environmental Research (BER) Program under Award Number DE-SC0018420) to EBC. Studies on fatty acid elongation were supported by the US DOE, Office of Science, BER project Bigger, Better Brassicaceae Biofuels and Bioproducts (B5) (Award Number DE-SC0023142) to EBC. Any opinions, findings, and conclusions or recommendations expressed in this publication are those of the author(s) and do not necessarily reflect the views of the U.S. Department of Energy. Research in the CZ lab was supported by the China Agriculture Research System (CARS-12). Molecular graphics and analyses performed with UCSF ChimeraX, developed by the Resource for Biocomputing, Visualization, and Informatics at the University of California, San Francisco, with support from National Institutes of Health R01-GM129325 and the Office of Cyber Infrastructure and Computational Biology, National Institute of Allergy and Infectious Diseases.

## Author Contributions

HK, KP, CZ, and EBC conceived the study. HK, KP, and EBC designed experiments and interpreted the data. HK, KP, and EBC wrote the manuscript. HK and KP generated expression vector constructs and managed transgenic plants. J-JMR assembled the PacBio Isoform Sequencing transcriptome and developed an internal BLAST server for homology searches. HK conducted gene expression analyses of *O. limprichtianus* and lipid analyses of transgenic *Arabidopsis*. KP conducted yeast experiments and Alphafold3 protein structure prediction. REC conducted TAG estolide analysis by LC-MS. HJ, FH, XT, TZ, and WY collected seeds and contributed to transcriptomic analyses and gene identification. EBC provided overall supervision for the project.

