## Supplementary Information for "Polyketide Synthase-Like Functionality Acquired by Plant Fatty Acid Elongase"

### Materials and Methods

#### Gene expression and gDNA PCR analyses

Total RNA from the young leaves, mature leaves, early developing seeds, and late developing seeds of *O. limprichtianus* was isolated using the RNeasy Plant Mini Kit according to the manufacturer's protocol (Qiagen). After the treatment of DNase I, total RNA (1 μg) was converted to cDNA using RevertAid first-strand cDNA synthesis kit with the oligo dT18 primer (Thermo Scientific). Genomic DNA was extracted from leaf of *O. limprichtianus* using CTAB method. Both Semi-quantitative RT-PCR and gDNA PCR were performed with Bio-Rad S1000 Thermal Cycler using the DreamTaq Green PCR Master Mix (Thermo Scientific). The primers used for PCR are listed in Supplementary table 1.

#### Protein 3D structure modeling

Protein sequences of OlFAE1-1, OlFAE1-2, AtKCR1, OlKCR1-1, and OlKCR1-2 were submitted to Google AlphaFold 3 server to predict protein structure (Abramson et al., 2024). To show ligand binding, NAD and oleic acids were included for structure prediction. Graphics and analysis were further processed using UCSF ChimeraX (Meng et al., 2023).

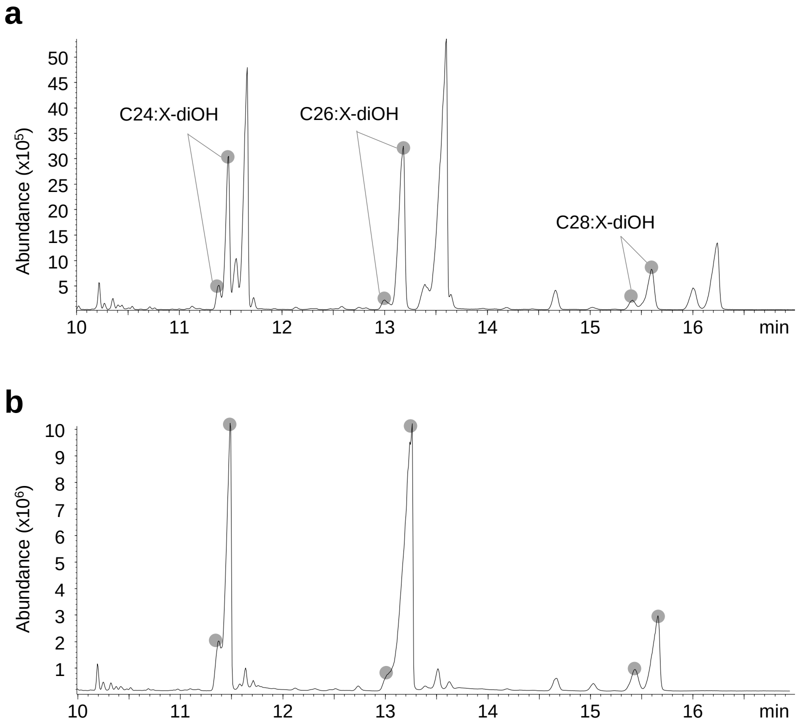


Supplementary Figure 1. Gas chromatography-mass analysis of trimethylsilyl (TMS) derivatives of FAMEs prepared from *O. limprichtianus* seeds.

**a**, TMS derivatives of FAMEs without reduction by sodium borohydride. **b**, TMS derivatives of FAMEs with reduction by sodium borohydride. The peaks pointed by grey circle represent derivatives of di-hydroxy fatty acids

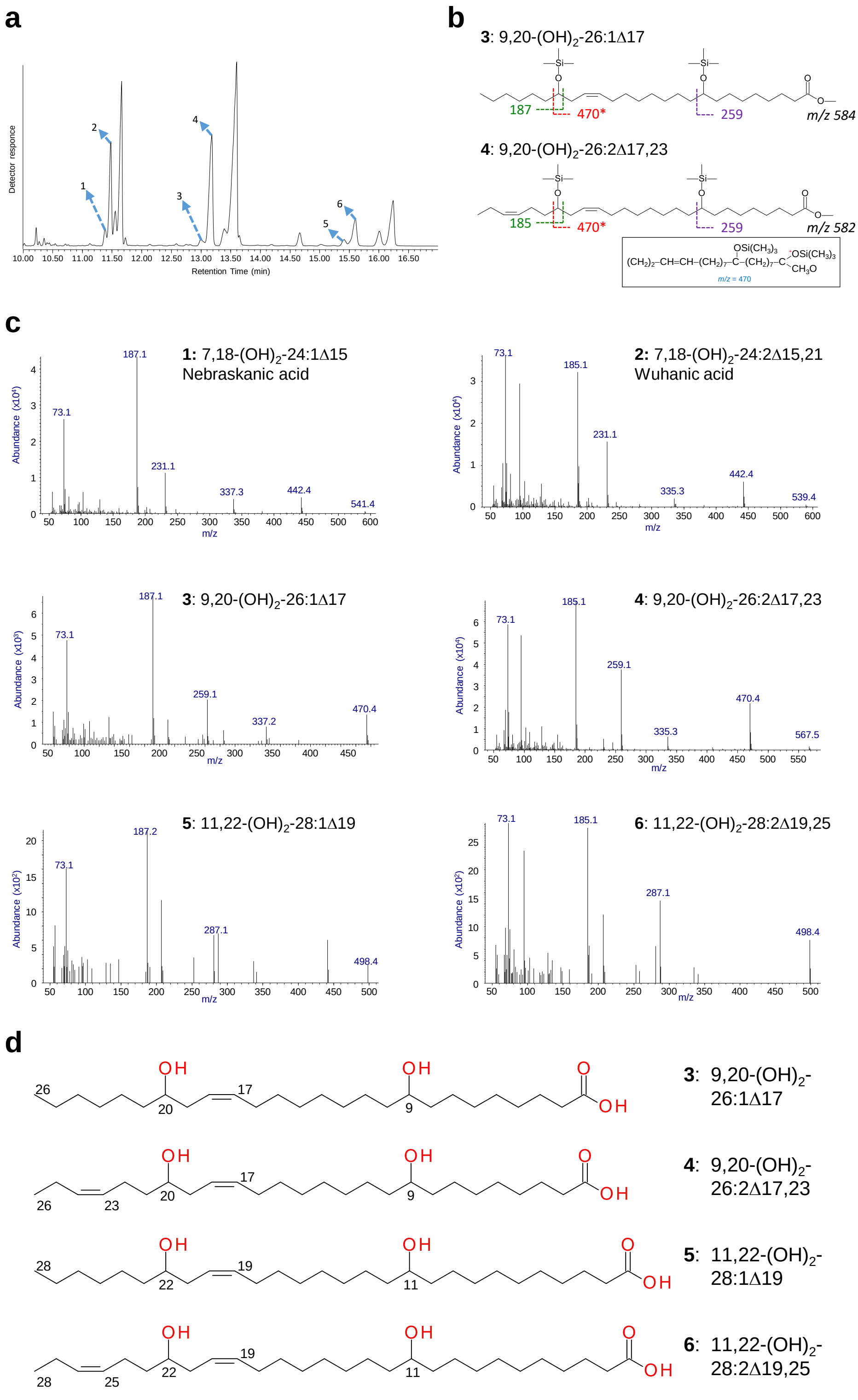


Supplementary Figure 2. GC-mass analysis of unusual fatty acids in *O. limprichtianus* seeds. **a**, GC-MS chromatogram trimethylsilyl (TMS) derivatives of FAMEs. **b**, Fragment ion patterns of peak 3 and 4. **c**, Mass spectra of peak 1 to 6. The asterisks for ion 'm/z = 470' in black box are formed by a complex rearrangement involving transfer of the TMS group to the carboxyl group. **d**, Structures of C26 and C28 di-hydroxy fatty acids.

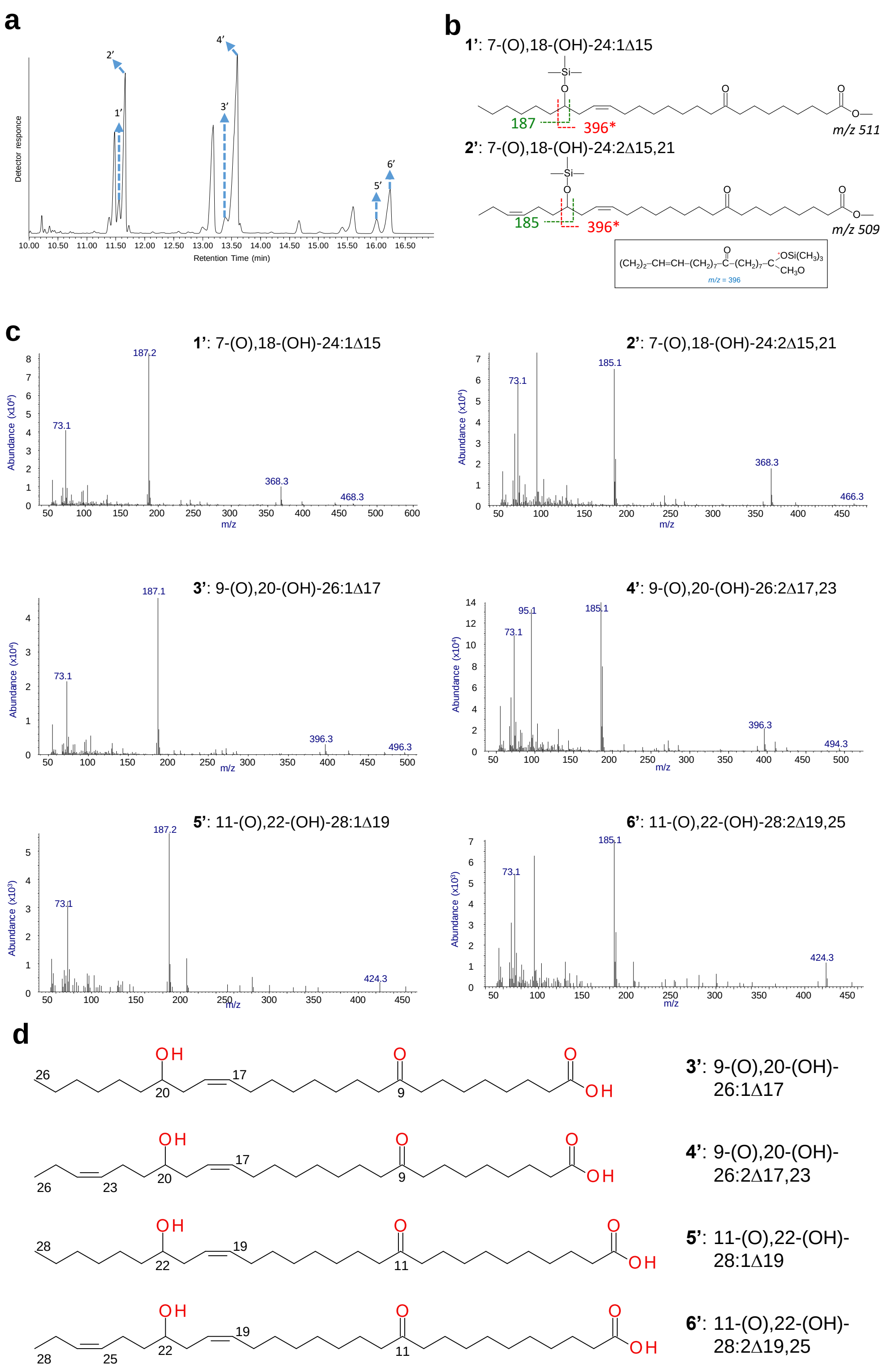


Supplementary Figure 3. GC-mass analysis of unusual fatty acids in *O. limprichtianus* seeds. **a**, GC-MS chromatogram trimethylsilyl (TMS) derivatives of FAMEs. **b**, Fragment ion patterns of peak 1' and 2'. **c**, Mass spectra of peak 1 to 6. The asterisk for ion 'm/z = 368' in is formed by a complex rearrangement involving transfer of the TMS group to the carboxyl group. **d**, Structures of C26 and C28 keto-hydroxy fatty acids.

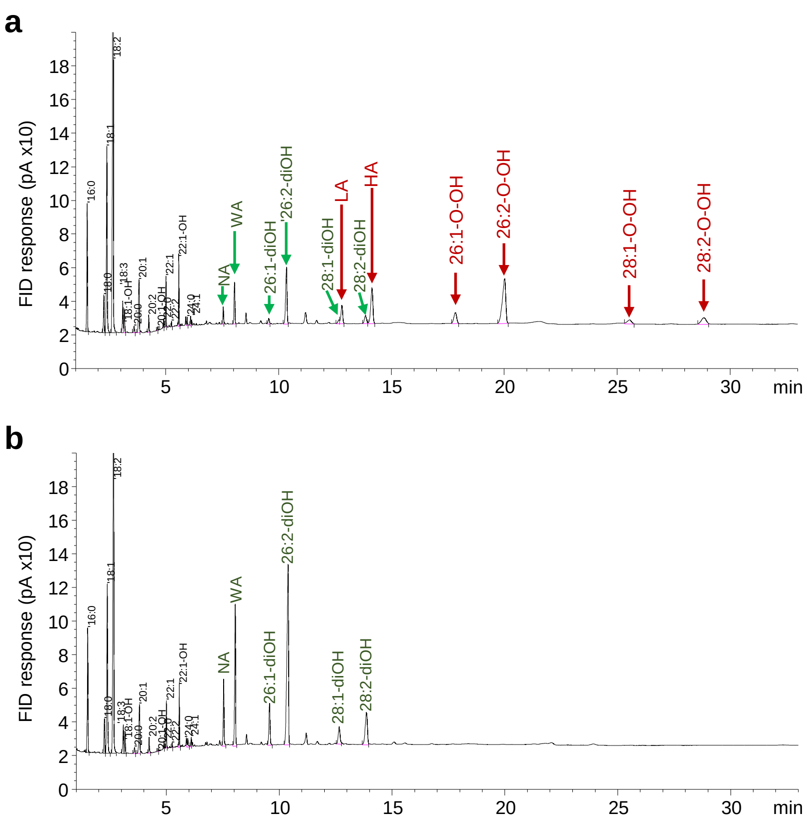



Supplementary Figure 4. Gas chromatography-FID analysis of trimethylsilyl (TMS) derivatives of FAMEs prepared from *O. limprichtianus* seeds.

**a**, TMS derivatives of FAMEs without reduction by sodium borohydride. **b**, TMS derivatives of FAMEs with reduction by sodium borohydride. NA, nebraskanic acid. WA, wuhanic acid. LA, licolnic acid. HA, hubeic acid.

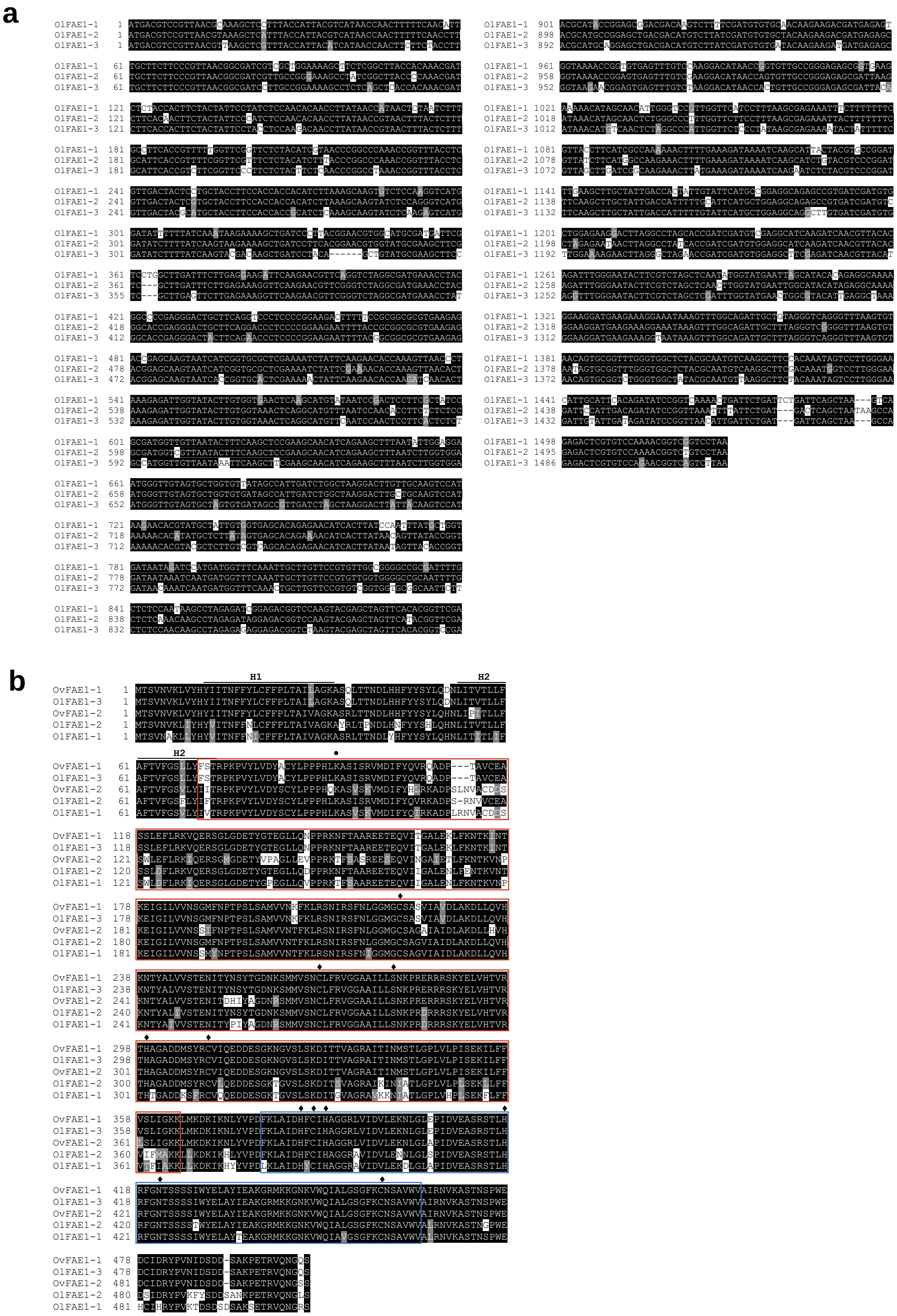


Supplementary Figure 5. The sequence analysis of FAE1 from Ol and Ov .

**a** and **b**, Boxshade analysis from aligned nucleotide (A), amino acid (B) sequecnes of *FAE1* genes from Ol and Ov. H1 and H2 indicate transmembrane domain. A close circle (•) indicate a residue Lys92 for catalytic activity and substrate specificity (Blocklock and Jaworski 2002). The acive sites are indicated by diamond (♦) (Millar et al. 1999). Red box indicates a domain for FAE1/Type III polyketide synthase-like protein. Blue box indicates a domain for 3-oxoacyl-(ACP) synthase III. Domain architecture analysis was performed using SMART version 9 (Letunic et al. 2021).

Blacklock, B. J., and Jaworski, J. G. (2002). Studies into factors contributing to substrate specificity of membrane-bound 3-ketoacyl-CoA synthases. European Journal of Biochemistry, 269(19), 4789-4798.

Millar, A. A., Clemens, S., Zachgo, S., Giblin, E. M., Taylor, D. C., and Kunst, L. (1999). CUT1, an Arabidopsis gene required for cuticular wax biosynthesis and pollen fertility, encodes a very-long-chain fatty acid condensing enzyme. The Plant Cell, 11(5), 825-838.

Letunic, Ivica, Supriya Khedkar, and Peer Bork. "SMART: recent updates, new developments and status in 2020." Nucleic acids research 49.D1 (2021): D458-D460.

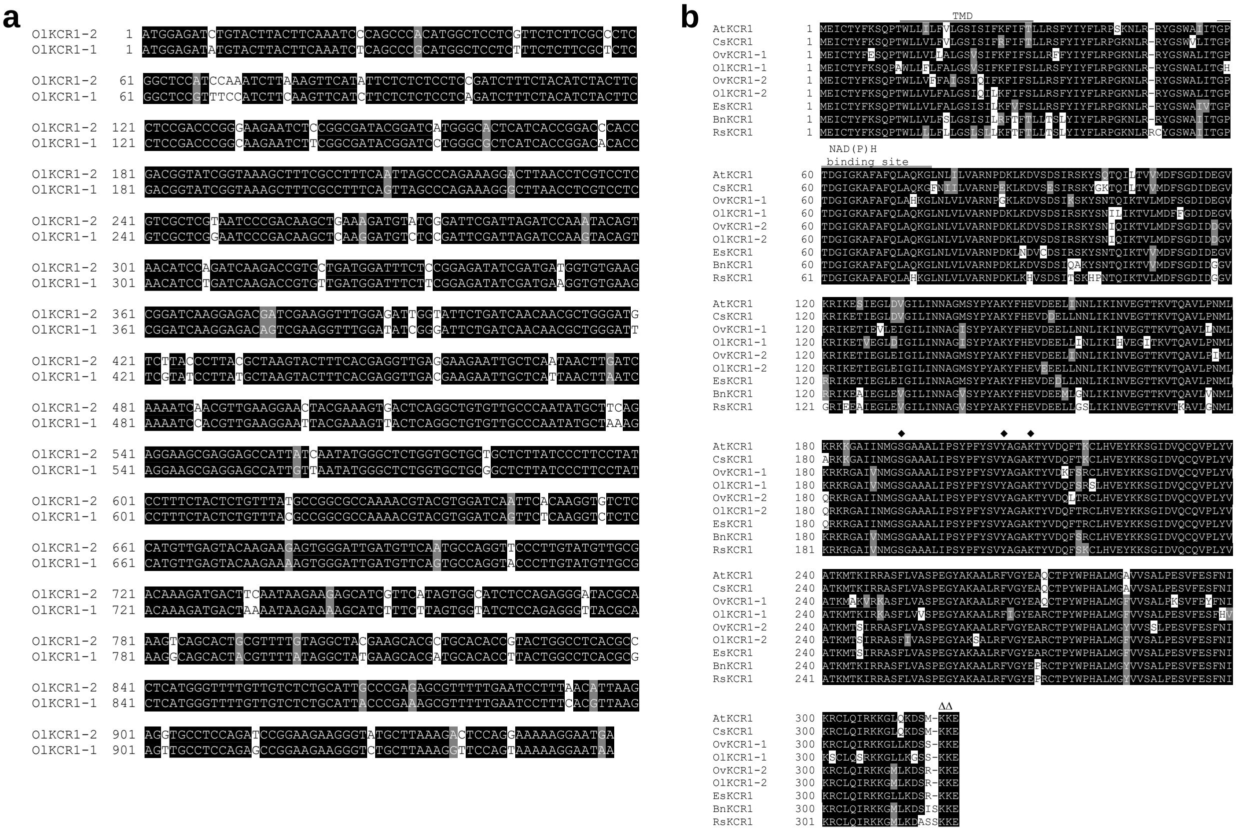


Supplementary Figure 6. Boxshade analysis from aligned DNA sequecnes of OlKCR1 genes (**a**) and aligned deduced amino acid sequences (**b**) of KCR1 genes from Ol, Ov (OvKCR1-1, OV09G031750 and OvKCR1-2, OV12G016450; http://www.bioinformaticslab.cn/pubs/OV_data/; Zhang et al. 2022), *Arabidopsis thaliana* (At; At1G67730), *Camelina sativa* (Cs; XP_010415354.1), *Brassica napus* (Bn; NP_001302805.1), *Raphanus sativus* (Rs; XP_018446636.2), and *Eutrema salsugineum* (Es; XP_006391228.1). TMD, transmembrane domain. The highly conserved serine (S), tyrosine (Y), and lysine (K) for the active triad are indicated by diamond (♦). The ER retention signal marked by triangle (∆).

Zhang, K., Yang, Y., Zhang, X., Zhang, L., Fu, Y., Guo, Z., ... & Cheng, F. (2023). The genome of Orychophragmus violaceus provides genomic insights into the evolution of Brassicaceae polyploidization and its distinct traits. Plant Communications, 4(2).

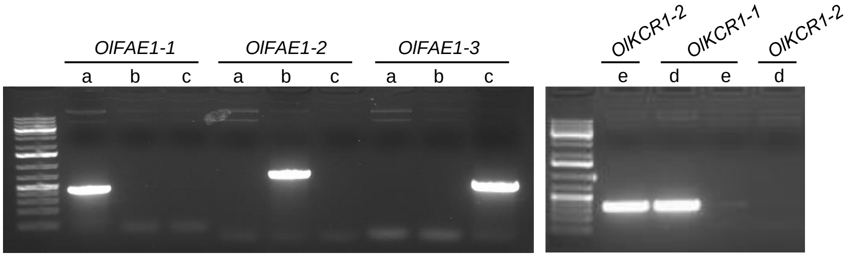


Supplementary Figure 7. PCR analysis for gene-specific primer-binding affinity. DNA framents were amplified using different plasmids as a DNA template. a, OlFAE1-1 in pBinGlyRed2 plasmid. b, OlFAE1-2 in pBinGlyRed2 plasmid. c, OlFAE1-3 in pBinGlyRed2 plasmid. d, OlKCR1-1 in pBinGlyRed2 plasmid. e, OlKCR1-2 in pBinGlyRed2 plasmid.

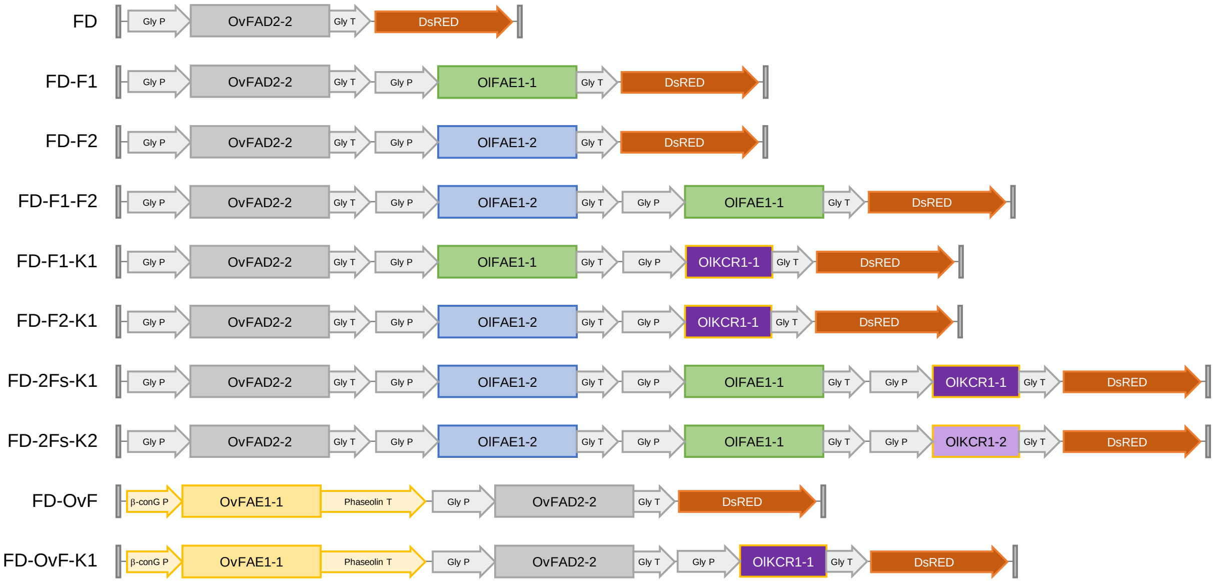


Supplementary Figure 8. Shematic diagram of T-DNA region of binary vectors used in this study. β-conG P, β-conglycinin promoter. Phaseolin T, phaseolin terminator. Gly P, glycinin promoter. Gly T, glycinin terminator.

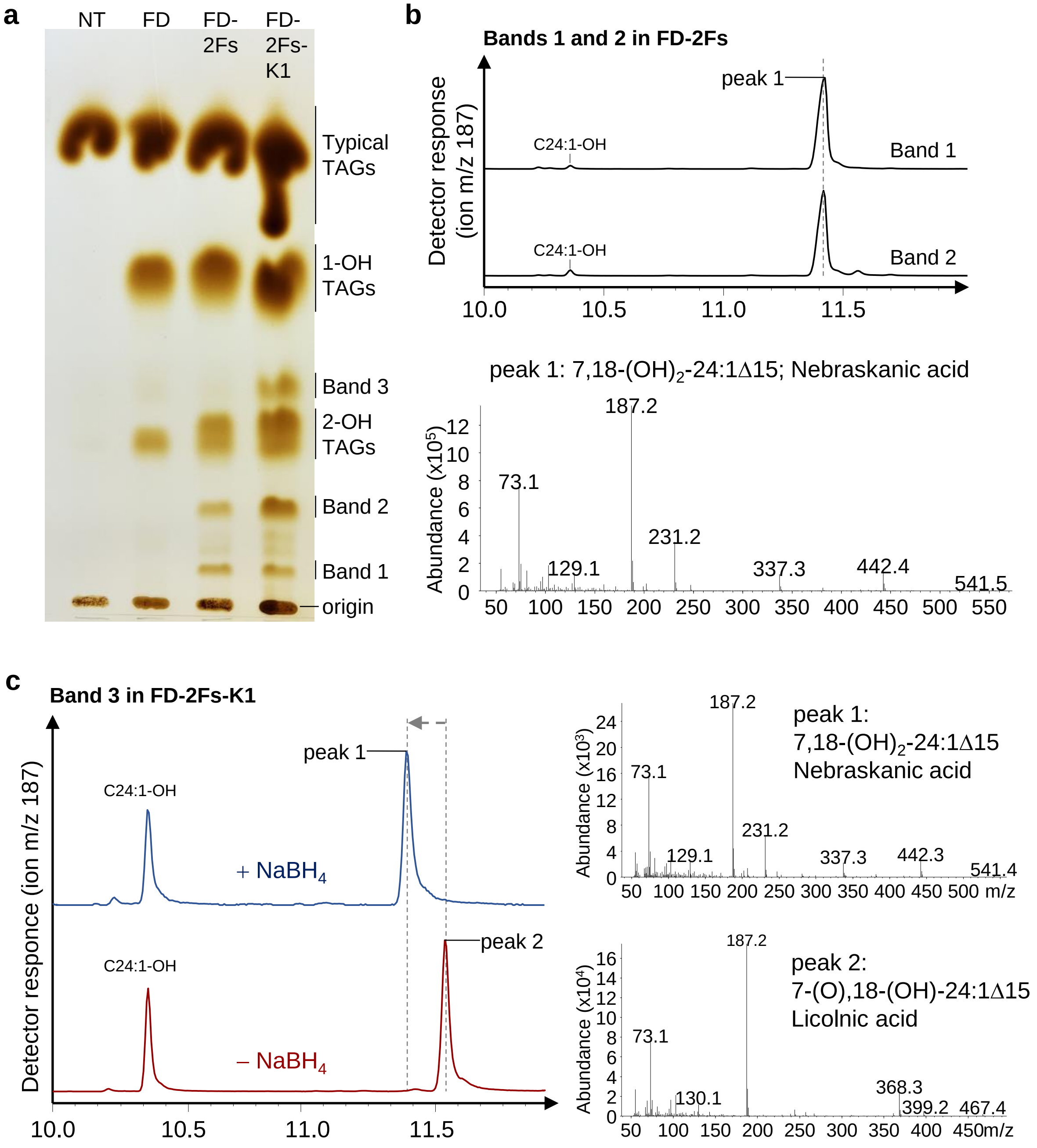


Supplementary Figure 9. Characterization of transgenic Arabidopsis oil composition.
**a**, TLC separation of major TAG classes from transgenic Col-0 seeds. 1-OH TAGs, TAG species containing one mono-hydroxy fatty acid. 2-OH TAGs, TAG species containing two mono-hydroxy fatty acid. **b** and **c**, GC-MS chromatograms and spectra for bands 1 and 2 of the FD-2Fs line (**b**) and band 3 of the FD-2Fs-K1 line (**c**). Bands 1 and 2 of the FD-2Fs line and band 3 of the FD-2Fs-K1 line were scrapped, converted to FAMEs, and then silylated for GC-MS analysis. Chromatograms display the extracted ion *m/z* 187 to identify the TMS derivatives of mono-unsaturated hydroxy fatty acids.

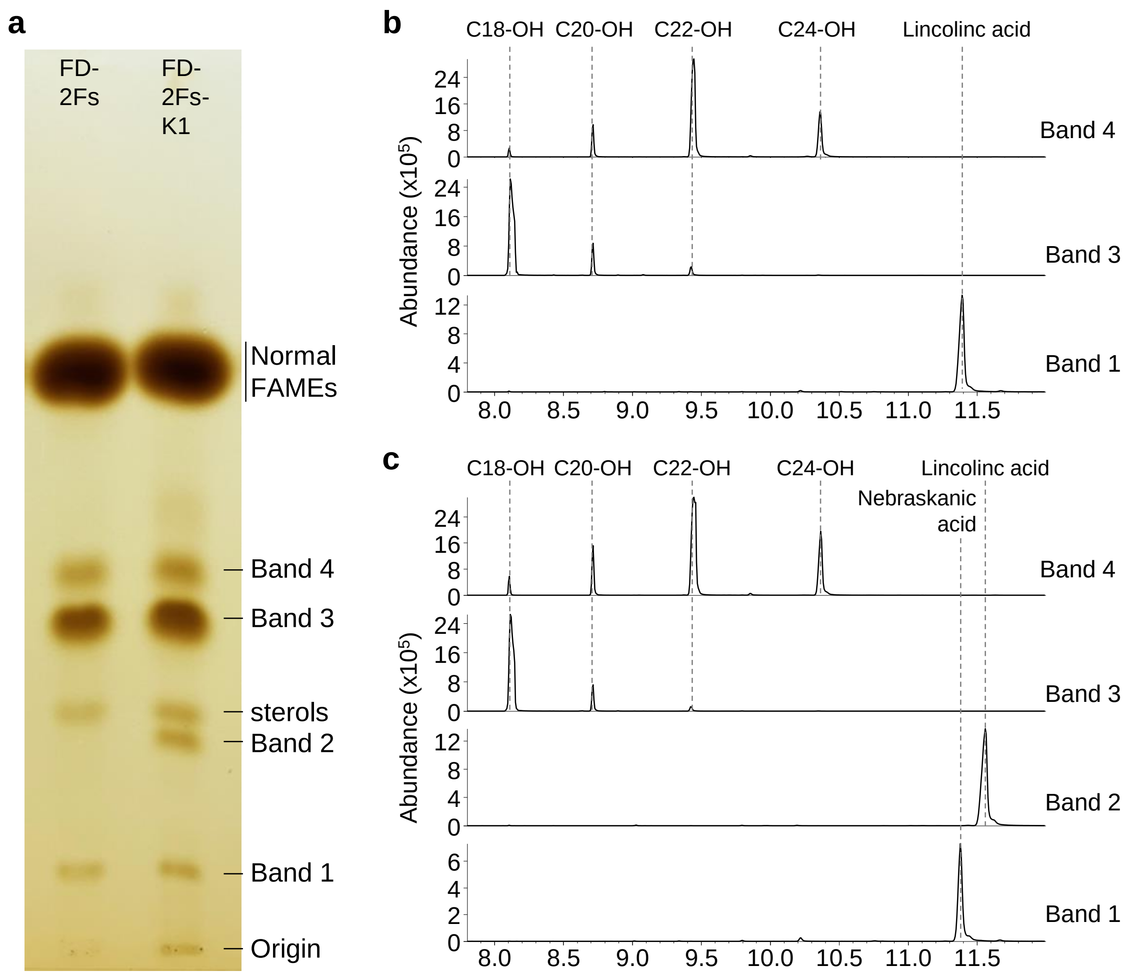


Supplementary Figure 10. Characterization of transgenic Arabidopsis fatty acid composition.
**a**, Total oils were converted to FAMEs and run TLC. **b** and **c**, GC-MS chromatograms showing the extracted ion m/z 187 to identify the TMS derivatives of mono-unsaturated hydroxy fatty acids from FD-2Fs line (**b**) and FD-2Fs-K1 line (**c**).

**a**

**b**

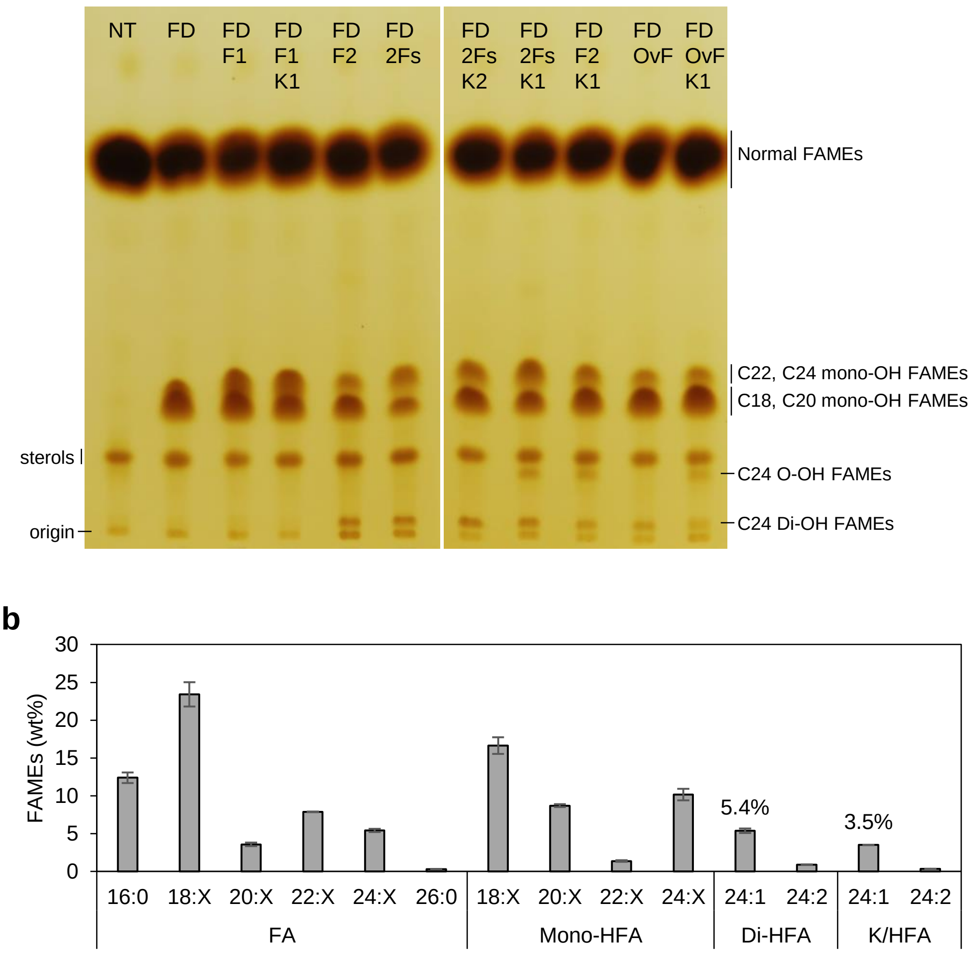


Supplementary Figure 11. Characterization of transgenic Col-0 seeds expressing various combination of *OlFAE1* genes and *OlKCR1* genes along with OvFAD2 hydroxylase.

**a**, FAMEs prepared from transgenic Arabidopsis seeds were separated on TLC plate with mobile phase [Diethyl Ether: Heptane; 60:40]. **b**, Fatty acid composition of FD-2Fs-K1 Arabidopsis seeds. Values are presented as wt% of total fatty acids. FA, fatty acids. Mono-HFA, monohydroxy fatty acids. Di-HFA, dihydroxy fatty acids. K/HFA, keto-hydroxy fatty acids. The results are the average of three replicates ± standard deviation.

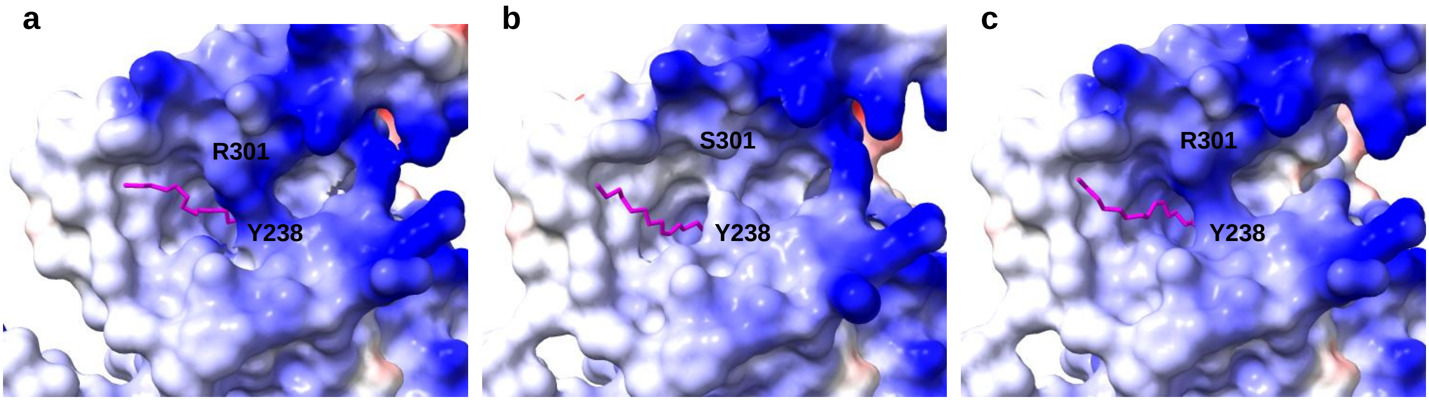


Supplementary Figure 12. Structure Prediction of KCR1 Proteins. Protein structures of KCR1 were predicted using A lphaFold 3 in the presence of NAD and oleic acid as ligands. (**a**-**c**) depict the structures of AtKCR1, OlKCR1-1, and OlKCR1-2, respectively. Electrostatic surface potentials are represented by color: white indicates neutral residues, bl ue indicates positive charges, and red indicates negative charges. Oleic acid molecules, shown in magenta, are loca lized within the FA binding pockets of the KCR proteins.

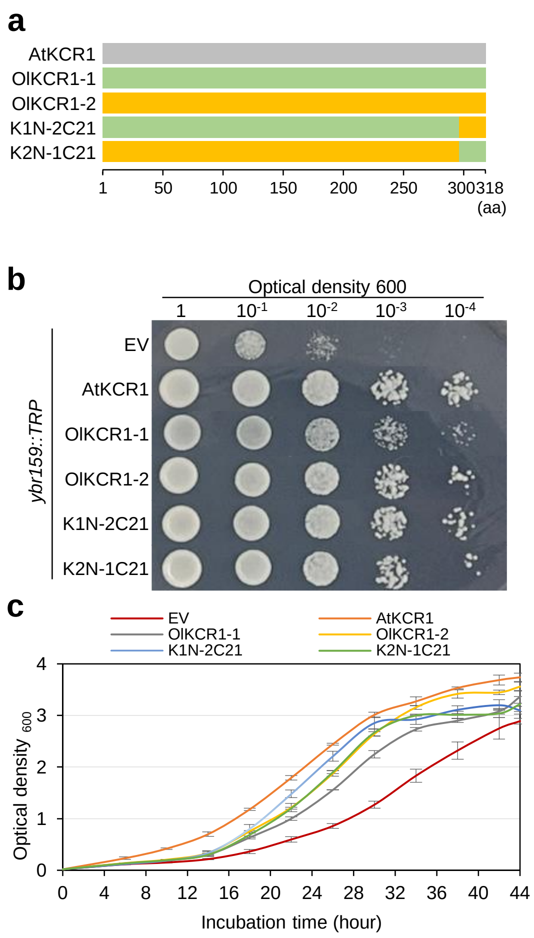


Supplementary Figure 13. Domain analysis of KCR1 in yeast.

**a**, Schematic view of native and chimeric KCR1 proteins. Domain swapped chimeric mutants were generated through overlapping PCR. K1N-2C21, OlKCR1-1$^{(1\text{-}297)}$ + OlKCR1-2$^{(298\text{-}318)}$. K2N-1C21, OlKCR1-2$^{(1\text{-}297)}$ + OlKCR1-1$^{(298\text{-}318)}$. **b** and **c**, Growth complement assay in *ybr159* yeast mutant. Vectors were introduced into *ybr159::TRP* strain. Cells harboring designated vector were inoculated under selective solid and liquid media (SD-Leu-Trp) at 30℃.

**a**

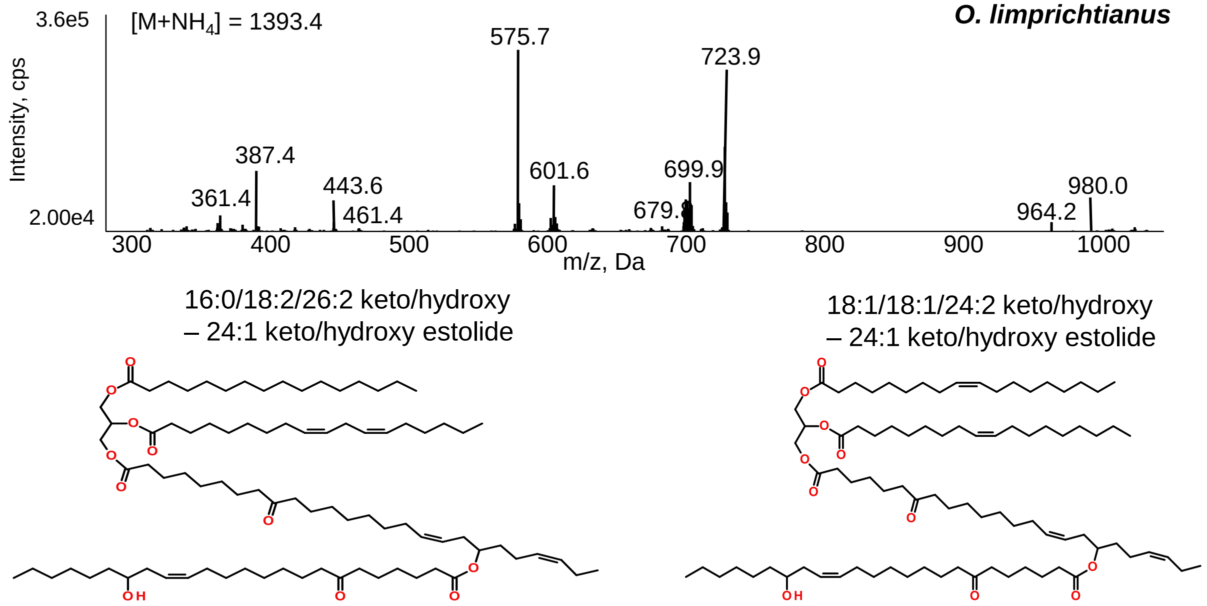


**b**

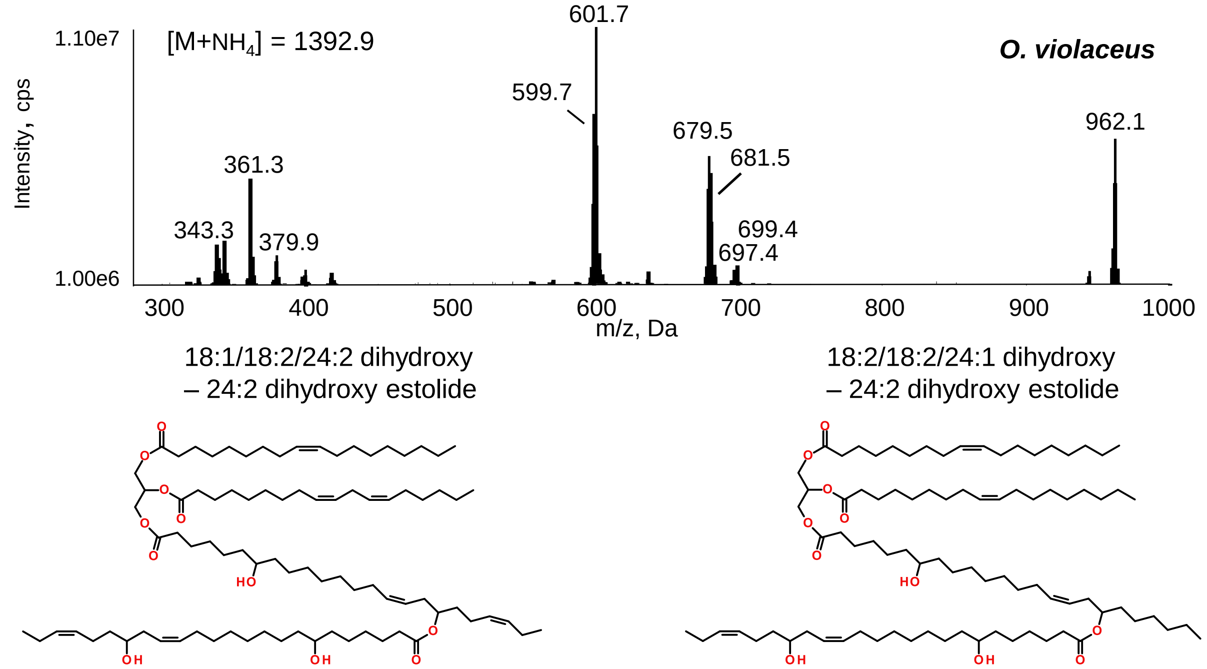


Supplementary Figure S14. Alternative estolide structures (m/z 1393) identified in Ol and Ov seed oils based on product ion spectra.
**a**, In Ol, two estolide species with identical precursor ions (m/z 1393.4 $[M+NH_4]^+$; $C_{87}H_{154}O_{11}$) were detected: 16:0/18:2/26:2 keto/hydroxy–24:1 keto/hydroxy estolide and 18:1/18:1/24:2 keto/hydroxy–24:1 keto/hydroxy estolide. Minor amounts of 16:0/18:2/26:2 dihydroxy–24:2 ketohydroxy estolide were also observed. Diagnostic product ions included 18:1/18:2 DAG (m/z 601.7), 18:2/18:2 DAG (m/z 599.7), 16:0/18:2 DAG (m/z 575.7), 18:2/26:2 keto-DAG (m/z 723.9; loss of 16:0), and 16:0/26:2 keto-DAG (m/z 699.9; loss of 18:2), the latter lacking the ability to lose a water molecule. **b**, In Ov, estolide species with m/z 1392.9–1393.2 $[M+NH_4]^+$ ($C_{87}H_{154}O_{11}$) were assigned as 18:1/18:2/24:2 dihydroxy–24:2 dihydroxy estolide or 18:2/18:2/24:1 dihydroxy–24:2 dihydroxy estolide. Corresponding product ions included 18:2/24:2 monohydroxy DAG (m/z 699.4), 18:2/24:3 monohydroxy DAG (m/z 697.4), and dehydrated derivatives at m/z 681.5 and 679.5, corresponding to 18:2/24:3 and 18:2/24:4 DAGs, respectively.

Supplementary Table 1. Primers used in this study.

| Primer list | Sequence 5'-3' | Explanation |
|---|---|---|
| o13F | ctcaagctcagcaccctacttc | Bind to Glycin promoter in pBinGlyRed2 vector |
| o14R | cgtgactcattatagagagtacttacat | Bind to Glycin terminator in pBinGlyRed2 vector |
| FAE1_F | CCGACACAAACAGAGCAATG | Isolation of FAE1 gene |
| FAE1_R | TCAGATACATCAAATTA | Isolation of FAE1 gene |
| KCR1_F | CCATTTCATCCATCTTCTCATG | Isolation of KCR1 gene |
| KCR1_R | CGTACAAGTCTTACAAAGAT | Isolation of KCR1 gene |
| OlFAE1-1-N-term_AS | GCACCTCTTAATGTTAAAGGATTCAAAAAC | paired with o13F, to generate chimeric 1 |
| OlFAE1-2-M-term_SE | GCACCTCTTAATGTTAAAGGATTCAAAAAC | paired with o14R, to generate chimeric 1 |
| OlFAE1-2-N-term_AS | GCAACTCTTAACGTGAAAGGATTCAAAAAC | paired with o13F, to generate chimeric 2 |
| OlFAE1-1-M-term_SE | GTTTTTGAATCCTTTCACGTTAAGAGTTGC | paired with o14R, to generate chimeric 2 |
| OlFAE1_C-term_SE | GAGATTGGTATACTTGTGGT | To generate Chimeric 3 and 4 |
| OlFAE1_C-term_AS | ACCACAAGTATACCAATCTC | To generate Chimeric 3 and 4 |
| OlKCR1-2C_5SE | AGCGTTTTTGAATCCTTTAACATTAAGAGGT | paired with o14R. to generate OlKCR1-1N + 1-2C |
| Chimeric1-1_3AS | GAGGCACCTCTTAATGTTAAAGGATTCAAAAACGCT | paired with o13F. to generate OlKCR1-1N + 1-2C |
| Chimeric1-1_5SE | AGCGTTTTTGAATCCTTTCACGTTAAGAGTTGCCTC | paired with o14R. to generate OlKCR1-2N + 1-1C |
| Chimeric1-2_3AS | GAGGCAACTCTTAACGTGAAAGGATTCAAAAACGCT | paired with o13F. to generate OlKCR1-2N + 1-1C |
| AtKCR1 5'-EcoRI | cggaattcATGGAGATCTGCACTTACTTCAAAT | To amplify AtKCR1 |
| AtKCR1 3'-XhoI | cgctcgagCCTGGAAGATTCATTCTTTCTTCATG | To amplify AtKCR1 |
| >H298N V299I OlKCR | cccgaaagcgtttttgaatcctttAacAttaagagttgcctccagagccgg | Primers for site-directed mutagenesis of OlKCR |
| >H298N V299I AS OlKCR | ccggctctggaggcaactcttaaTgtTaaaggattcaaaaacgctttcggg | Primers for site-directed mutagenesis of OlKCR |
| >S301R S305I SE OlKCR | cctttCacGttaagagAtgcctccagaTccggaagaagggtctgc | Primers for site-directed mutagenesis of OlKCR |
| >S301R S305I AS OlKCR | gcagacccttcttccggAtctggaggcaTctcttaaCgtGaaagg | Primers for site-directed mutagenesis of OlKCR |
| >N298H I299V SE OlKCR | GAGCGTTTTTGAATCCTTTcACgTTAAGAGGTGCCTCCAGATC | Primers for site-directed mutagenesis of OlKCR |
| >N298H I299V AS OlKCR | GATCTGGAGGCACCTCTTAAcGTgAAAGGATTCAAAAACGCTC | Primers for site-directed mutagenesis of OlKCR |
| >R301S I305S SE OlKCR | CCTTTaACaTTAAGAGcTGCCTCCAGAgCCGGAAGAAGGGTATGCT | Primers for site-directed mutagenesis of OlKCR |

| | | |
|---|---|---|
| >R301S I305S AS OlKCR | AGCATACCCTTCTTCCGGcTCTGGAGGCAgCTCTTAAtGTtAAAGG | Primers for site-directed mutagenesis of OlKCR |
| >G313D SE OlKCR1-1 | gaagaagggtctgcttaaagAttccagtaaaaaggaataac | Primers for site-directed mutagenesis of OlKCR |
| >G313D AS OlKCR1-1 | gttattcctttttactggaaTctttaagcagacccttcttc | Primers for site-directed mutagenesis of OlKCR |
| >D313G SE OlKCR1-2 | GAAGAAGGGTATGCTTAAAGgCTCCAGGAAAAAGGAATGACTCG | Primers for site-directed mutagenesis of OlKCR |
| >D313G AS OlKCR1-2 | CGAGTCATTCCTTTTTCCTGGAGcCTTTAAGCATACCCTTCTTC | Primers for site-directed mutagenesis of OlKCR |
